# *Cryptococcus neoformans* resists to drastic conditions by switching to viable but non-culturable cell phenotype

**DOI:** 10.1101/552836

**Authors:** Benjamin Hommel, Aude Sturny-Leclère, Stevenn Volant, Nathanaël Veluppillai, Magalie Duchateau, Chen-Hsin Yu, Véronique Hourdel, Hugo Varet, Mariette Matondo, John R Perfect, Arturo Casadevall, Françoise Dromer, Alexandre Alanio

## Abstract

Metabolically quiescent pathogens can persist in a viable non-replicating state for months or even years. For certain infectious diseases, such as tuberculosis, cryptococcosis, histoplasmosis, latent infection is a corollary of this dormant state, which has the risk for reactivation and clinical disease. During murine cryptococcosis and macrophage uptake, stress and host immunity induce *Cryptococcus neoformans* heterogeneity with the generation of a sub-population of yeasts that manifests a phenotype compatible with dormancy (low stress response, latency of growth). In this subpopulation, mitochondrial transcriptional activity is regulated and this phenotype has been considered as a hallmark of quiescence in stem cells. Based on these findings, we worked to reproduce this phenotype in vitro and then standardize the experimental conditions to consistently generate this dormancy in *C. neoformans*.

We found that incubation of stationary phase yeasts (STAT) in nutriment limited conditions and hypoxia for 8 days (8D-HYPOx) was able to produced cells that mimic the phenotype obtained *in vivo*. In these conditions, mortality and/or apoptosis occurred in less than 5% of the yeasts compared to 30-40% of apoptotic or dead yeasts upon incubation in normoxia (8D-NORMOx). Yeasts in 8D-HYPOx harbored a lower stress response, delayed growth and less that 1% of culturability on agar plates, suggesting that these yeasts are viable but non culturable cells (VBNC). These VBNC were able to reactivate in the presence of pantothenic acid, a vitamin that is known to be involved in quorum sensing and a precursor of acetyl-CoA. Global metabolism of 8D-HYPOx cells showed some specific requirements and was globally shut down compared to 8D-NORMOx and STAT conditions. Mitochondrial analyses showed that the mitochondrial masse increased with mitochondria mostly depolarized in 8D-HYPOx compared to 8D-NORMox, with increased expression of mitochondrial genes.

Proteomic and transcriptomic analyses of 8D-HYPOx revealed that the number of secreted proteins and transcripts detected also decreased compared to 8D-NORMOx and STAT, and the proteome, secretome and transcriptome harbored specific profiles that are engaged as soon as four days of incubation. Importantly, acetyl-CoA and the fatty acid pathway involving mitochondria are required for the generation and viability maintenance of VBNC.

Altogether, these data show that we were able to generate for the first time VBNC phenotype in *C. neoformans*. This VBNC state is associated with a specific metabolism that should be further studied to understand dormancy/quiescence in this yeast.

**Authors Summary:** Quiescence/dormancy in microorganism is a common feature that enables survival at the population level. In fungi, quiescence has been studied in the baker yeast *Saccharomyces cerevisiae*. In *Cryptococcus neoformans*, dormancy is of great interest since it is known from the natural history of cryptococcosis that dormancy in yeast can last decades exists before possible reactivation upon immunosuppression. Based on a previous study which identified a subpopulation of dormant yeasts in experimental models of cryptococcosis, we identified here in vitro conditions that enabled the induction of dormancy via the formation of viable but non culturable cells (VBNC). Reactivation of part of these cells was possible through stimulation with vitamin B5, a quorum sensing molecule. We showed that the global metabolism of the VBNC was down but harbored specific signatures compared to control conditions. We identified mitochondrial metabolism, in particular the fatty acid pathway, as key for the maintenance and viability of VNBC. These findings open the road for research on dormancy. Elucidating the parameters involved will help understand the pathophysiology of the disease including the difficulty in eradication of the yeasts despite therapy, and the possible relapse/recurrence of the infection.

## Introduction

All microorganisms are exposed to fluctuating environments and periodic external stresses, which inhibit their growth. Some microbes may overcome these stress periods by entering into a resistant non-replicative state, assessed by /associated with a decreased culturability [1]. This decrease was first assumed to reflect microbial death as a consequence of a stochastic and induced nutrient deprivation decay [2]. In quantum mechanics, the Schrödinger’s experiment considered a cat simultaneously alive and dead until a measurement was made. A parallel experiment could be done with a dormant microorganism. Answering if it is alive or dead seems trivial but is unquestionably complex [3]. In fact, there are two other alternative explanations. Non-culturable cells could be the resulting state of a genetically programmed death or could be viable but non culturable cells (VBNC) that are adapted to living / surviving as dormant cells either by the formation of spores (bacterial of fungal) or by modification of the metabolic state of the initial cell [4,5].

There is now clear evidence that VBNC are viable cells with an intact cell membrane and a low metabolic activity [6]. Various environmental stressors can induce VBNC, such as starvation, hypoxia, stressful temperature, pH variations [7]. In a finalistic approach, to enable perpetuation of the lineage, VBNC should be capable at some point to exit the dormant state to resume an active and replicating form when environmental conditions improve. This process could be described as reactivation or “resuscitation”. At least 85 species of bacteria [8] and 10 species of fungi [9–11] have been reported capable of adapting a VBNC state. Hence, the VBNC state seems to be an intrinsic capability of many microorganisms including pathogenic ones [12–15]. During infection, the “dormant” pathogens are able to tolerate both immune system attacks and prolonged exposure to antimicrobials [16]. Pathogens are not known to cause disease when present in the VBNC state but virulence is retained and infection can be initiated again following reactivation [6].

Dormancy is an important pathogenic feature for some invasive fungal diseases, such as cryptococcosis [17–19] and histoplasmosis [12], where dormant organisms are known to reactivate years after primary infection. Cryptococcosis is caused by the ubiquitous environmental basidiomycete yeast *Cryptococcus neoformans*. It occurs mainly in immunocompromised individuals, and especially those with AIDS, with more than 200 000 new cases of meningoencephalitis and more than 180, 000 deaths per year worldwide [20]. Latency is an important phase of cryptococcosis pathogenesis. However, the biological features of the primary cryptococcal infection in humans has been rarely investigated [21], in contrast to tuberculosis. Inhalation of infectious propagules leads to a primary pulmonary infection with a granulomatous immune response [21,22], that either restrains infection or eradicates the yeast. This immune response can fail when immunosuppression occurs leading to reactivation and disease.

Recently, evidence was reported for the existence of dormant yeast cells of *C. neoformans* in models of *C. neoformans*/host interaction *in vivo* (mice) and *in vitro* (macrophages) [23]. A population of dormant yeast cells harboring characteristics that could be related to dormancy (low metabolic response, growth latency, increased mitochondrial activity, increased autophagy, decreased neoglucogenesis and reactivation by fetal calf serum) was described [23]. To characterize these cells, we needed reproducible *in vitro* conditions that would generate yeast cells with a phenotype identical to that observed in the subpopulations of yeasts upon interaction with hosts (decreased stress response, delay of growth). Hypoxia and nutrient starvation are the two main factors that are known to allow induction of dormancy in various cell types (including mammalian stem cells, and bacteria). Based on previous studies performed on dormancy of skeletal muscle stem cells, it is known that hypoxia is able to induce and maintain dormancy [24]. To induce quiescent *M. tuberculosis*, two well-known models are used: the Wayne model based on oxygen depletion in nutrient rich medium and the Loebel model based on nutrient deprivation in oxygen rich medium [25].

To induce dormancy in *C. neoformans*, we developed and standardized an *in vitro* model using two consecutives stresses, i.e. nutrient starvation and hypoxia. We then analyzed the parameters influencing dormancy and reactivation of dormant yeasts. Using phenotypic microarrays along with proteome, secretome, and transcriptome analyses, we further validated and characterized these yeast cells and the metabolic pathways that are crucial for the generation, maintenance of and exit from dormancy.

## Results

### Yeasts incubated in hypoxia were intact and exhibited low stress response and delayed growth

Based on our previous identification of dormant cells in a mouse model of cryptococcosis [23], and on conditions (nutrient and oxygen limitation) used to study quiescence in *M. tuberculosis* [25] we identified *in vitro* conditions that induced a phenotype in *C. neoformans* cells that mimicked the phenotype observed *in vivo* (mainly decreased stress response, delay of growth). Yeasts grown in liquid YPD medium until stationary phase (STAT) which results in a drastic nutrients deprivation, were incubated without change/replenishment of medium (to avoid any additional nutrient stress) at 30°C for 8 days in hypoxia (O_2_<0.1%, these cells will be called 8D-HYPOx from now on) or normoxia (8D-NORMOx) (Supplementary Figure 1). The influence of prolonged incubation time (25 days), increased (37°C) or decreased (4°C) temperature of incubation, in the presence of hypoxia or normoxia was also tested during preliminary experiments but did not result in an interesting or reliable phenotype compared to the established protocol.

We first checked the impact of the selected conditions on the morphology and basic features of *C. neoformans* cells. Median cell diameters were not different for yeasts in 8D-HYPOx, 8D-NORMOx compared to STAT (4.5 µm; interquartile range (IQR) [3.8-5.1], 4.6 µm [4.1-5.0] and 4.6 µm [4.1-5.0], respectively). Median capsule thickness was smaller in 8D-HYPOx (0.8 µm [0.7-1.0] and 8D-NORMOx (0.9 µm [0.7-1.0]) compared to STAT (1.0 µm [0.9-1.2]), (Supplementary Figure 2A and B, p<0.0001). Specific staining revealed no difference in the shape of the nucleus (Figure 1A), nor in the capsule structure based on the anti-glucuronoxylomannan monoclonal antibody (E1) binding (Figure 1A, Supplementary Figure 3A). Staining with MDY-64 showed an increased quantity of vacuolar membrane (3.1 ± 0.4 increase in the geometric mean fluorescence for 8D-HYPOx/8D-NORMOx compared to STAT (Figure 1A, Supplementary Figure 3B). Plasma membrane was intact for the majority of cells (88% of 8D-HYPOx, 60% of 8D-NORMOx compared to 98% of STAT, Figure 1A, Supplementary Figure 3C). Transmission electron microscopy (TEM) revealed a significantly thicker cell wall in 8D-HYPOx compared to STAT (median 206 nm [170.5-245.0] vs. 163.5 nm [123.8-199.3], p<0.05, Supplementary Figure 2C) and the presence of large intracytoplasmic vacuoles (Figure 1B).

**Figure 1:**
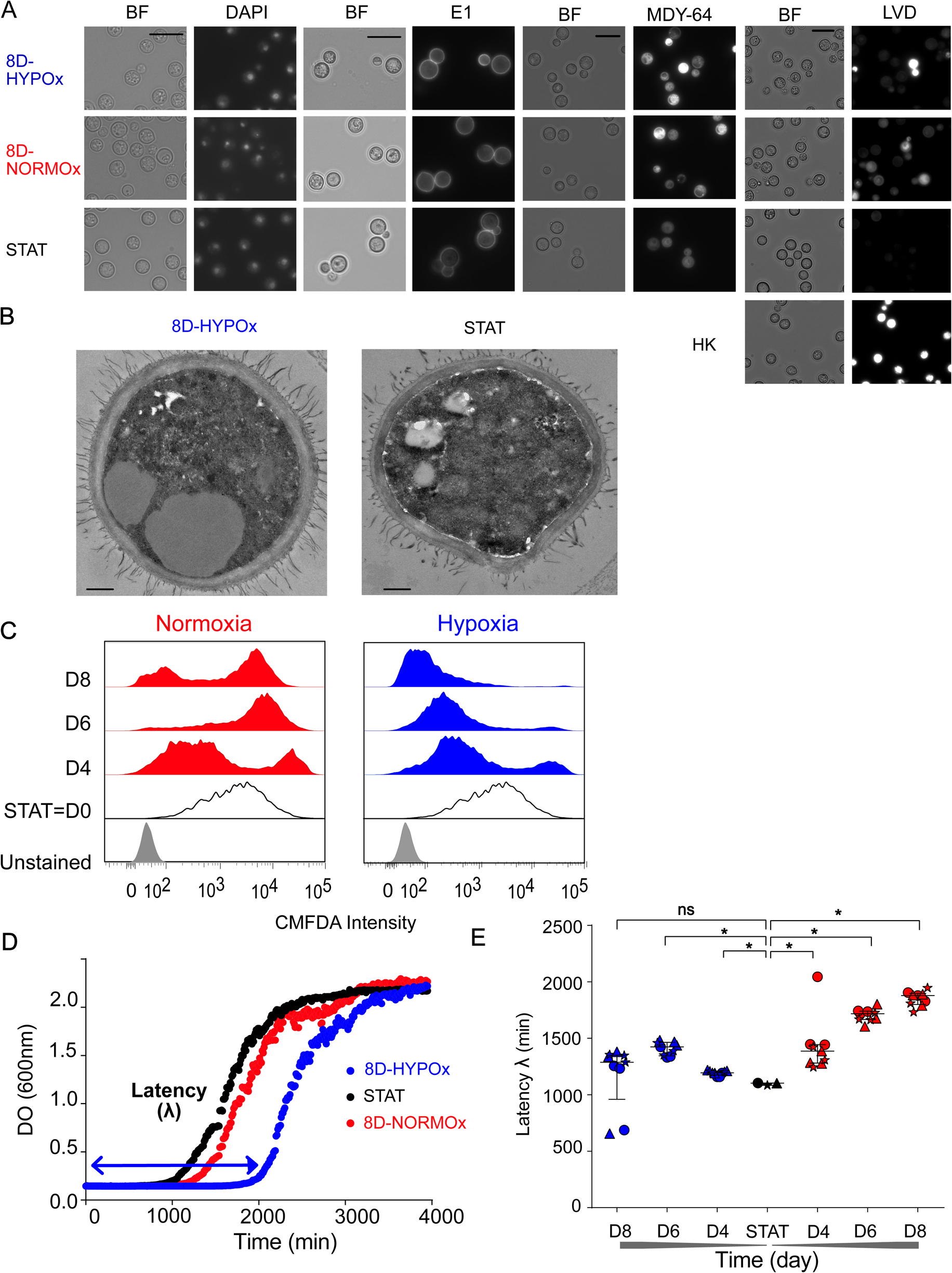
*Cryptococcus. neoformans* incubated for 8 days in hypoxia were intact with a low stress response and delayed growth. **A**. Cells recovered after 8 days in hypoxia (8D-HYPOx) were compared to those incubated in normoxia (8D-NORMOx), to stationary phase (STAT=D0) cells. DAPI staining (DAPI) showed intact cells with a nucleus; labeling with the E1 anti-glucuronoxylomannan monoclonal antibody showed similar staining pattern on the cryptococcal capsule in all conditions; vacuolar staining MDY-64 showing more intense staining in 8D-HYPOx/8D-NORMOx than STAT yeasts; LiveDead^®^ violet staining showing high viability in 8D-HYPOx (HK = heat killed yeasts, BF = bright field, bar = 10µm) **B**. Transmission electron microscopy of 8D-HYPOx and STAT showing the presence of large intracytoplasmic vacuoles (bar = 0.5 µm) **C**. CMFDA labeling showed that 8D-HYPOx yeasts had decreased stress response over time compared to 8D-NORMOx. Representative from over 5 independent experiments. **D**. Representative growth curves in the different conditions with 8D-HYPOx showing an increased latency (λ) as calculated upon mathematical modelling. **E**. Latency of growth increased over time for 8D-HYPOx compared to STAT and 8D-NORMOx (*p <0.01, ns not significant). Experiments were done 3 times independently (each experiment is identified with a different symbol, circle, triangle, star). Each replicate represents the value of the latency (λ) for each biological replicate as the mean of 3 technical replicates. Medians and interquartile range (IQR) are represented.

We then verified if the stress response and delay in growth were observed as *in vivo* [23]. Intracellular levels of glutathione, a well-established marker of stress response in various mammalian cell types [26], fungi [27] including *C. neoformans* [23], were assessed kinetically by using 5-chloromethylfluorescein diacetate (CMFDA) staining at day 0 (STAT) and up to day 8 (D4, D6, D8). A decrease in stress response was observed over time with an apparently homogenous population harboring a low CMFDA intensity in 8D-HYPOx, as opposed to 8D-NORMOx for which a double population (low and high CMFDA intensity corresponding to low and high stress response) was observed (Figure 1C). As a lag phase was observed in “dormant” cells *in vivo*, we then studied the growth curves upon incubation in rich liquid YPD (growth curve method) for 8D-HYPOx, 8D-NORMOx and STAT and compared them after curve fitting using latency (lag phase λ) as a major readout (Figure 1D). Median latency was significantly increased for 8D-NORMOx and 8D-HYPOx compared to STAT (1103 min [1083-1104]. It increased over time for 8D-HYPOx from 1387 min [1281-1445] at D4 to 1878 min [1800-1898] at D8, whereas it plateaued for 8D-NORMOx (1194 min [1176-1210] at D4, and 1288 min [960-1351] at D8) (Figure 1E).

Since latency of growth can be influenced by numerous factors including cell concentration and culture medium (representing the metabolic potential of the cells influenced by the quantity of nutrients), we tested these parameters. Serial 10-fold dilutions of the yeasts (3×10^5^ to 3/well) resulted in an increased latency for both 8D-HYPOx and STAT, in undiluted YPD and minimal medium (MM) (Supplementary Figure 4A-B). The latency was higher in MM than in YPD for 8D-HYPOx at all the dilutions studied (Table 1) and for STAT at the lowest dilutions (<3×10^3^/well). Low nutrient medium (MM) increased the latency by a differential of 2505 min for STAT and 2656 min for 8D-HYPOx compared to a richer medium (YPD). When extrapolating the linear regression for 1 cell we observed only little difference between latency in 8D-HYPOx and STAT (Δ8D-HYPOx-STAT) for YPD (1820 min) or for MM (1971 min). The slope of the growth curve that represents the rate of cell generation during the exponential phase was comparable for 8D-HYPOx, 8D-NORMOx, and STAT.

**Table 1:**
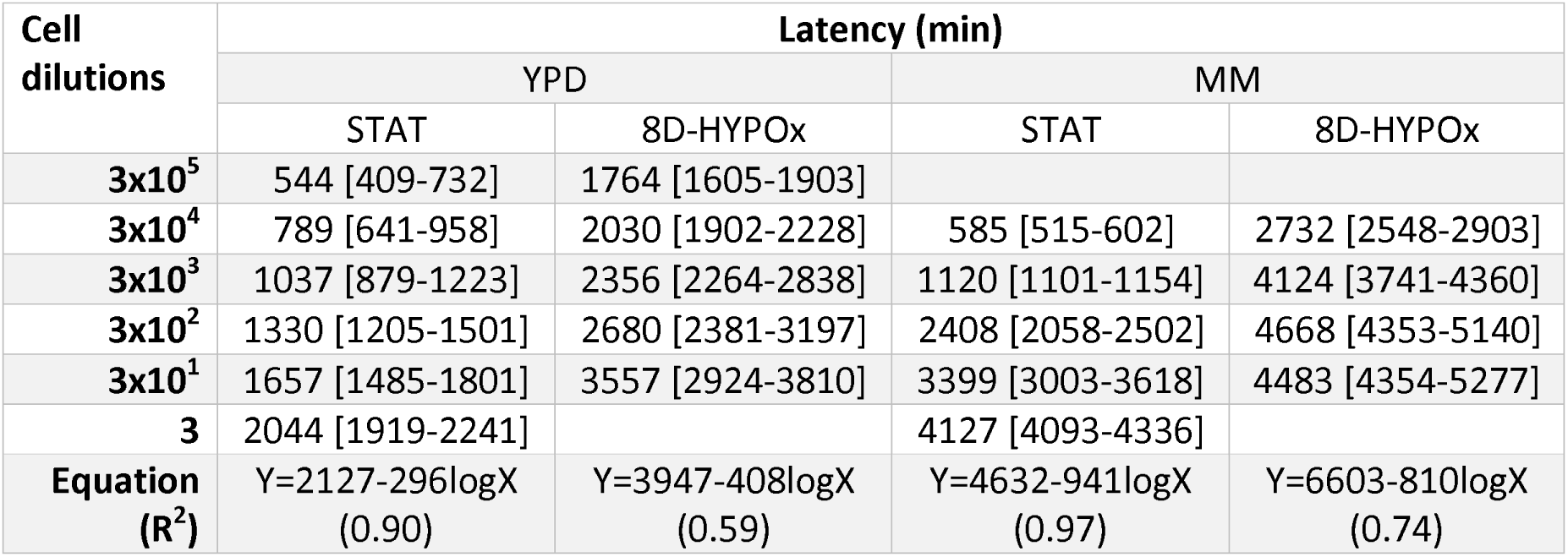
Influence of cell dilutions and culture medium on the latency obtain from the growth curves.

### Yeasts incubated in hypoxia were mostly viable but non-culturable cells (VBNC)

We then investigated the viability of the yeasts incubated in hypoxia based on nucleic acids stains and apoptosis assays. Cell viability and apoptosis-like phenomenon were measured over time and evolved differently in hypoxia and normoxia (Figure 2A). Viable non-apoptotic cells represented 98.7% [98.4-99.0]) in the STAT condition. In hypoxia, the proportion decreased at D2, D4 and D6, culminating at 74.6% [70.5-78.8] at D4 and reaching 98.9% [98.8-99.0]) in 8D-HYPOx. In normoxia, the proportion decreased and plateaued at D8 at 63.6% [62.2-65.0]. The proportion of apoptotic-like cells, negligible in STAT cells (0.8% [0.5-1.6]), peaked at D4 in hypoxia (21.4% [17.5-25.4]) while it increased over time to reach a plateau at D6 in normoxia (35.8% [34.9-36.7]). Dead cells represented less than 3% over time in hypoxia (2.7% [1.7-3.7] at D4), while it reached 5% [4.2-5.9] at D6 in normoxia and was less than 1 % in STAT. In parallel, culturability was assessed based on the colony forming unit method (CFU method) by plating on solid medium (YPD agar). From 100% in STAT, the culturability decreased in both conditions but significantly more in 8D-HYPOx than in 8D-NORMOx (17.5% [15.4-19.7], vs. 0.8% [0.5-1.0]; p<0.01) (Figure 2B). Since a small proportion of cells were cultivable by the CFU method, we wondered if this explained the delay in growth observed by the growth curve method (Figure 1D-1E). Indeed, 8D-HYPOx included a small subpopulation of cells harboring a high CMFDA staining (high stress response) (Figure 2C). We thus hypothesized that they were the metabolically active cells responsible for the observed growth, and that they should differ from the other subpopulations. 8D-HYPOx subpopulations were sorted according to their CMFDA intensity (low, medium and high). The latency of the three subpopulations was comparable to that of the unsorted (bulk) population and significantly higher than that of the 8D-NORMOx and STAT (Figure 2C). Likewise, culturability assessed by the CFU method was not significantly different in CMFDA^low^ (0.19±0.20), CMFDA^medium^ (0.28±0.17), CMFDA^high^ (0.10±0.11) compared to the unsorted population (0.53±0.41) (p>0.01), although significantly lower than for 8D-NORMOx (12.73% [7.62-18.62]) and STAT (102.5% [93.25-111.8]) (Figure 2C, p<0.01). We then assess by a limited dilution method (plating method) based on the Poisson law the culturability of the different subpopulations seeded at 3333 cells/well in liquid YPD in 96-well plates. In this assay, the number of positive wells / number seeded allows comparison of the culturability. A decrease in culturability was found for 8D-HYPOx with 10 /18 (55.5%) for the CMFDA^low^, 16/18 (88.8%) for the CMFDA^medium^, 14/18 (77.7%) for the CMFDA^high^, and 17/18 (94.4%) for the unsorted populations whereas all wells (100%) grew for the 8D-NORMOx (6/6) and STAT (3/3) (Supplementary Figure 4C).

**Figure 2:**
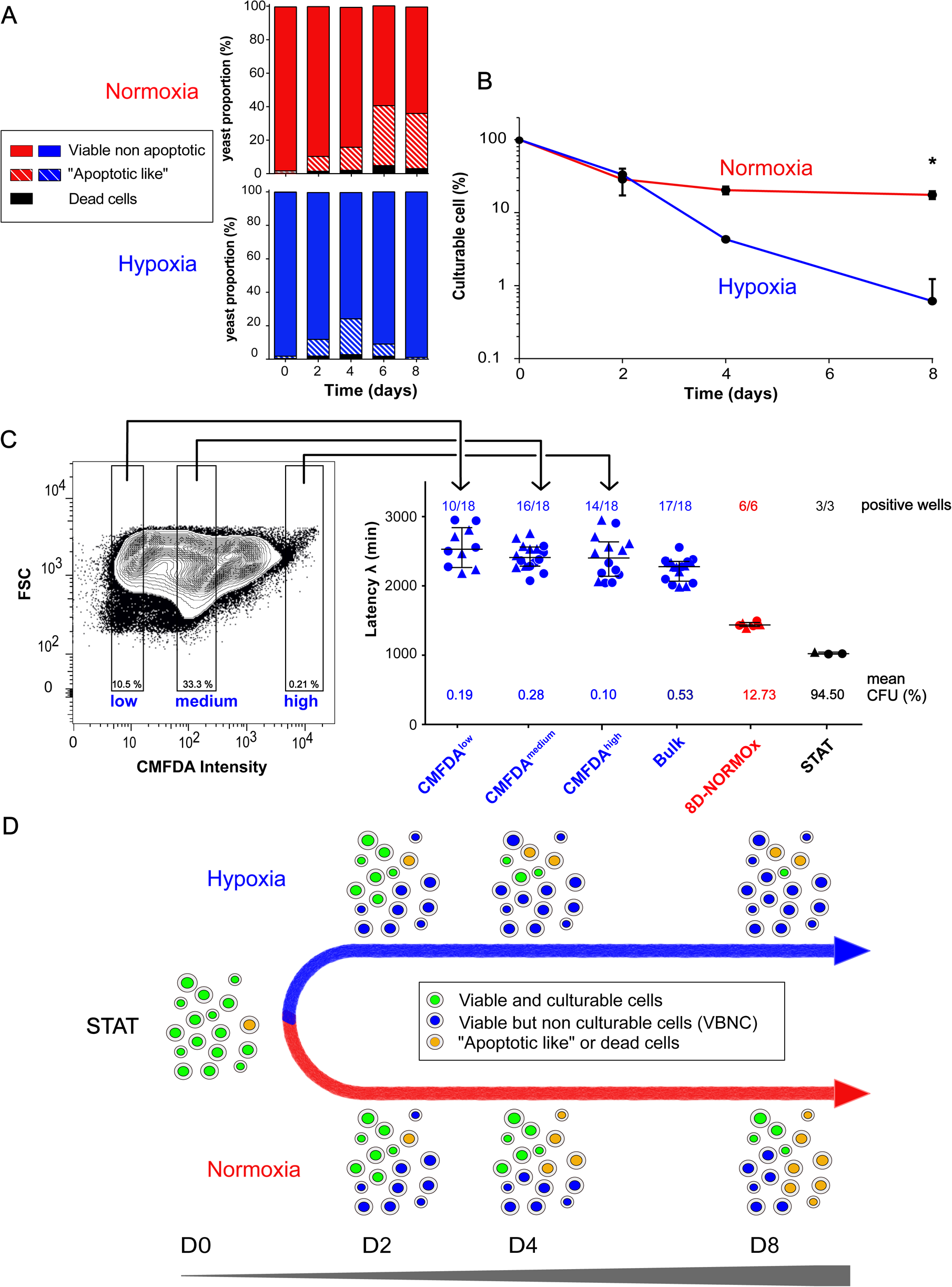
*Cryptococcus neoformans* incubated for 8 days in hypoxia harbor a homogenous phenotype of viable but non-culturable cell (VBNC). **A.** Kinetics analysis using multispectral flow cytometry with viability (LIVE/DEAD^®^) and apoptosis (TUNEL) staining showing a homogenous population of viable yeasts after D8 in hypoxia and a heterogeneous population in normoxia with a mixture of viable, apoptotic like and dead yeasts. The experiment was done in triplicate with the median value represented. Viable non-apoptotic cells represented a median proportion of 87.7% [84.2-91.2] at D2, 74.6% [70.5-78.8] at D4, 90.9% [89.7-92.2] at D6, and 98.9% [98.8-99.0] at D8 in hypoxia, 89.4% [88.4-90.5] at D2, 83.5% [83.1-83.9] at D4, 59.6% [59.2-60.0] at D6, 63.6% [62.2-65.0] at D8 in normoxia and 98.7% [98.4-99.0] in STAT. Apoptotic-like cells represented a median proportion 9.9% [6.2-13.8] at D2, 21.4% [17.5-25.4] at D4, 7.2% [6.5-7.9] at D6, 1.1% [0.9-1.2] at D8 in hypoxia, 8.8% [7.7-10.0] at D2, 13.8% [13.6-14] at D4, 35.8% [34.9-36.7] at D6, 32.9% [31.0-34.9] at D8 in normoxia and 0.8% [0.5-1.6] in STAT cells. The dead cells median proportion was 0.5% [0.4-0.6] in STAT, 1.9% [1.8-2.0] at D2, 2.7% [1.7-3.7] at D4, 1.8% [1.6-2.0] at D6, 0% at D8 in hypoxia1.7% [1.6-1.8] at D2, 2.2% [1.8-2.6] at D4, 5.0% [4.2-5.9] at D6, 3.2% [3.0-3.4] at D8 in normoxia. **B.** Culturability was followed over 8 days by plating yeasts on Sabouraud agar (colony forming unit method). The proportion of culturable cells decreased significantly over time under hypoxia and significantly more than in 8D-NORMOx. Compared to STAT condition (110.5% [108-113]), the median culturability was (33.2% [31.1-35.2] at D2, 4.3% [4.0 −4.6] at D4, 0.8% [0.5-1.0] at D8) in hypoxia and (28.6% [17.2-40.0] at D2, 20.3% [17.9-22.8] at D4 and 17.5% [15.4-19.7] at D8 in normoxia cells) (* p<0.01). **C.** Cell sorting according to the CMFDA intensity allowed showing a latency of growth after 8D-HYPOx and was increased regardless of the stress response and comparable in the three populations (low, medium, high) to that of the unsorted population (bulk) compared to 8D-NORMOx and STAT: CMFDA^low^ (2520 min [2257-2831]), CMFDA^medium^ (2401 min [2276-2553]) and CMFDA^high^ (2394 min [2129-2627]) was not significantly different from unsorted cells (2269 min [2059-2344]) (p>0.01) although significantly different from 8D-NORMOx (1427 min [1407-1462]) and STAT (1013 min [1107-1036]) conditions (* p<0.01). Two independent experiments were pooled (each experiment is identified by a different symbol, each dot represents a technical replicate). Mean culturability (% mean CFU after plating) reported below the plots was comparable in all hypoxia populations and lower than in the other conditions. The ratio of positive wells/seeded wells are reported above the plots for each population recovered at the end of the growth curve measurement. **D.** Proposed model for the evolution of yeasts phenotypes upon incubation in hypoxia or normoxia leading to the generation of a VBNC phenotype after 8D-HYPOx.

Overall, using different methods to assess culturability on solid (CFU method) and liquid medium (growth curves and plating method), only a small proportion of yeasts incubated in hypoxia and nutrient depletion were cultivable, and they were not distinguishable from the other subpopulations based on stress response. These results suggested that the dormancy induction conditions led to the production of a population of yeasts mostly composed of what is called viable and non-culturable cells (VBNC). A model for the evolution of the cell population phenotypes over time in hypoxia and normoxia is proposed in Figure 2D.

### 8D-HYPOx can be reactivated by pantothenic acid (PA), the acetyl CoA precursor

One of the major characteristics of VBNC is that they can be reactivated under specific conditions which should translate into a decrease in latency. To prove that the 8D-HYPOx were VBNC, we investigated factors that enabled their reactivation, i.e. switch from non-culturable to culturable cells, taking into account that a small proportion of the cells from the 8D-HYPOx bulk were already culturable.

We hypothesized that the latency observed in 8D-HYPOx could be impacted by the high quantity of nutrients available in the medium that would inhibit reactivation of yeasts previously adapted to starvation and thus unable to cope with high nutrient concentration in a new medium. If this was true, then incubation of 8D-HYPOx in diluted medium should have the reverse effect and result in decreased latency. This hypothesis was tested by refeeding 8D-HYPOx in MM diluted in water. MM dilutions resulted in a significant decrease in 8D-HYPOx latency (Figure 3A) from 4463 min [3947-5488] in 100%MM to 3722 min [3612-4156] in 20%MM and 3358 min [3240-3664] in 10%MM (p<0.01, Figure 3A). This decrease was also observed for STAT but to a lesser extent (1593 min [1397-1720], 1326 min [1220-1510] and 1245 min [1045-1277] in 100%MM 20%MM and 10%MM, respectively, p <0.01). Dilutions of nutrients was thus beneficial for yeasts growth.

**Figure 3:**
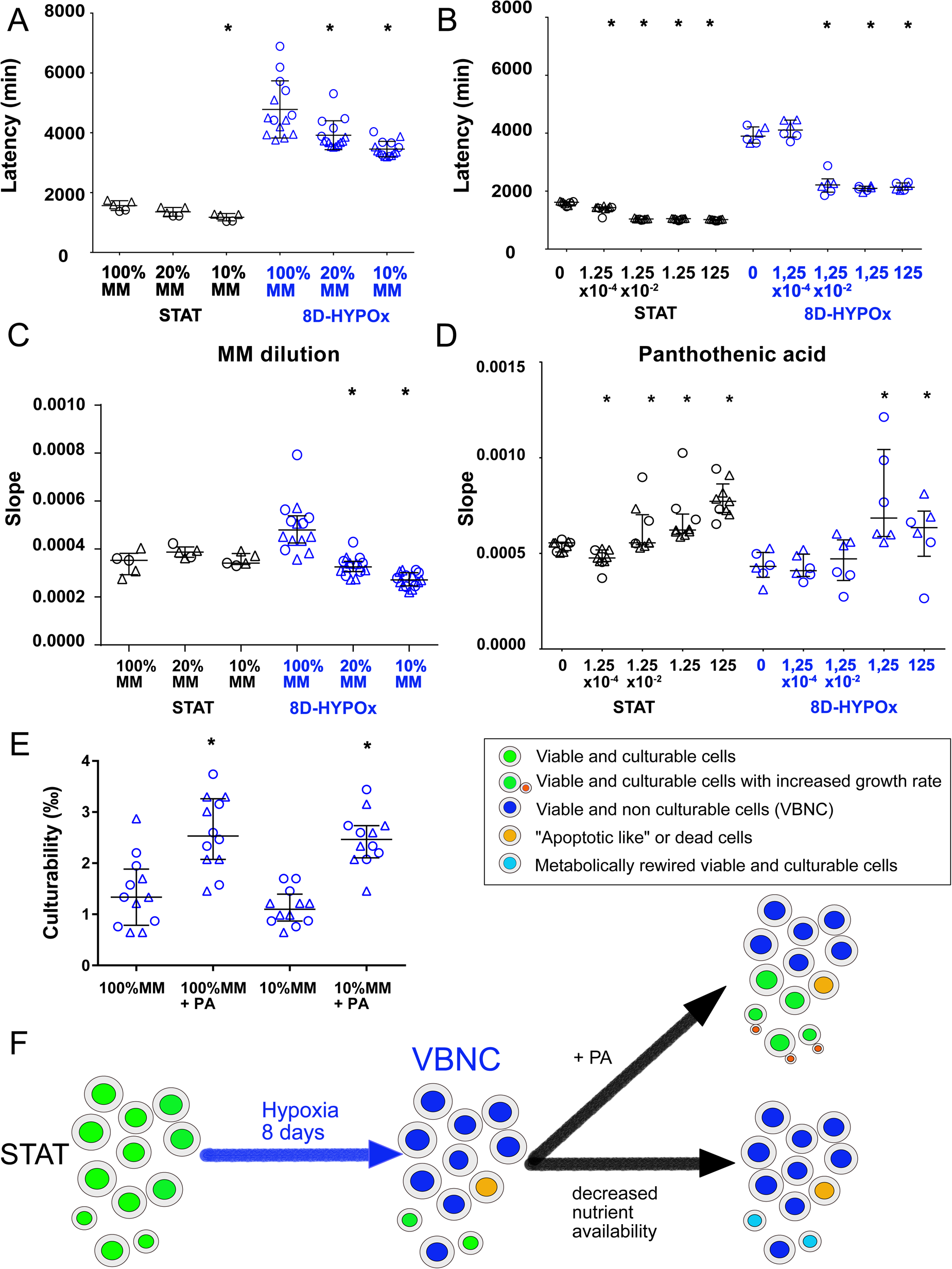
*Cryptococcus neoformans* incubated for 8 days in hypoxia reactivated with pantothenic acid (PA) and culturable cells rewired in minimal medium (MM) dilutions. Growth curves were determined on yeasts recovered as STAT or 8D-HYPOx and then refed in diluted MM. Latency and growth rate (slope) were then determined. Experiments were all repeated twice (independent experiments are identified as circle or triangle). Graph includes dot for the replicates, as well as median ± IQR (*p<0.01 compared to control in the same condition). **A.** The dilution of minimal medium from 100% to 10% (diluted MM) resulted in a progressive decrease of latency in 8D-HYPOx cells and to a lesser extend in STAT cells. (*p<0.01 compared to 100%MM). Two independent experiments were pooled. Each point represents the latency/well in two (STAT n=5, 8D-HYPOx n=15) **B.** 100-fold dilution of pantothenic acid (starting at 125 μM) were added to the suspension prior to growth curve determination. Pantothenic acid significantly decreased the latency at a concentration of 1.25×10^-2^ μM or more in 8D-HYPOx and to a lesser extend in STAT cells (* p<0.01). Two independent experiments were pooled. Each point represents the latency/well in two (STAT n=9, 8D-HYPOx n=6). **C.** Growth rate was significantly decreased in 20%MM and 10%MM in 8D-HYPOx **D.** A significantly increased growth rate was observed after addition of PA (1.25 µM or more) to 8D-HYPOx cells and at all concentrations for STAT. **E.** The addition of pantothenic acid at 125µM but not the dilution of MM increased the culturability of 8D-HYPOx based on the observation of the number of positive wells (100 cells per well) and the calculation of the probability of one cell to grow. Each dot represents the calculated culturability. Two independents experiments were pooled. **F.** Proposed model for the effect of hypoxia and nutrient depletion. From now on, 8D-HYPOx are called VBNC (Viable but not culturable cells). After 8 d of incubation in hypoxia and nutrient depletion, the decrease in latency observed upon addition of pantothenic acid on VBNC is explained by the reactivation of VBNC to culturable cells (increased culturability) and by the increased growth rate of culturable cells (increased slope). The decrease in latency observed upon dilutions of MM (decreased nutrient availability) is explained by a faster rewire of the culturable cells (decreased latency corresponding to decreased time to resume growth) without increased culturability or growth rate.

Then, we tested the effect of conditioned medium (freeze-dried and rehydrated MM recovered from STAT), murine macrophage cell (J774 cell line) lysate, and known cofactor of growth in yeasts (pantothenic acid (PA, vitamin B5), thiamine (vitamin B1), sodium pyruvate, and L-DOPA) on 8D-HYPOx growth. Latency decreased only upon addition of J774 lysate, conditioned medium and PA. As conditioned medium and J774 lysate are each complex media rich in PA, we decided to focus specifically on the effect of PA, based on the fact that PA was known to decrease latency in *C. neoformans* cells [28] and was a known as a secreted quorum sensing compound by *C. neoformans* [29]. Addition of PA to 100%MM showed a dramatic effect on the latency of 8D-HYPOx. For instance, latency was decreased at concentrations of PA above 1.25 x 10^-2^ μM from 3994 min [3797-4332] (no PA) and 4111 min [3866-4451] (1.25 x 10^-4^ µM) to 2213 min [1963-2426] (1.25 x 10^-2^ µM), 2095 min [2007-2170] (1.25 µM) and 2133 min [2035-2281] (125 µM) (Figure 3B, p <0.01). In comparison, less effect of PA was observed on STAT cells (from 1615 min [1532-1637] (no PA) to 1433 min [1398-1456] (1.25 x 10^-4^µM) and then 1036 min [1017-1045] (1.25 x 10^-2^µM), 1048 min [1017-1045] (1.25 µM), 1018 min [982.5-1031] (125 µM)).

Decreased in latency following addition of PA could result from increased growth rate or/and increased number of culturable yeasts and thus reactivation of VBNC. MM dilutions (20%MM and 10%MM) significantly decreased the growth rate (slope) for 8D-HYPOx compared to 100%MM but did not alter that of STAT cells (Figure 3C). PA significantly increased growth rate of 8D-HYPOx at concentrations above 1.25 µM and that of STAT cells at concentrations above 1.25 x 10^-2^µM (Figure 3D, p < 0.01).

We then assessed whether the number of culturable cells increased or whether 8D-HYPOx exhibited improved metabolic adaptation. We used the plating method to calculate the probability for one cell to grow (Supplementary Figure 4C) after addition of PA or MM dilution. Dilution of MM alone (10%MM) did not modify the median proportion of culturable cells, but addition of PA increased it approximately two-fold from 1.3 ‰ [0.8-1.9] to 2.5 ‰ [2.0-3.3] in 100%MM and from 1.1 ‰ [0.9-1, 4] to 2.5 ‰ [2.1-2.7] in 10%MM. (p <0.01, Figure 3E). We also wondered whether phagocytosis would change culturability which was not the case with a comparable median proportion of culturable cells (1.04 ‰ [0.78-1.48]) and after 2 h macrophage uptake (1.21 ‰ [1.09-1.33], p=0.35; Supplementary Figure 4D).

Overall, when decreasing nutrient availability (diluting medium), latency decreased without change in the number of culturable yeast cells, suggesting that culturable yeasts were better adapted to grow in poor than in richer medium. By contrast, addition of PA allowed VBNC to switch to culturable cells (cell reactivation) with a decreased latency reflected an increasing number of culturable cells. However, the 2-fold increase in culturability after addition of PA was not sufficient to explain the extent of the decrease in latency suggesting that PA influenced also growth adaptation (Figure 3F). Altogether, 8D-HYPOx yeast harbored characteristics of VBNC and will be called VBNC from now on.

We then explored the characteristics of the VBNC as these cells have never been described in *C. neoformans* and rarely in other fungi, except for *Saccharomyces cerevisiae*. We explored the physiologic substrate required by VBNC using the Biolog^®^ phenotypic microarray, as well as their mitochondrial status, their secretome, proteome and transcriptome.

### VBNC showed specific metabolic requirements and low global metabolic activity

The physiologic substrates required for the metabolism of VBNC were analyzed by phenotypic microarrays using Biolog^®^ substrate utilization tests and compared with that of yeast cells in other conditions. Based on the 761 metabolites tested and excluding 240 compounds positive in the dead cells control, hierarchical clustering showed that VBNC possessed a unique profile (Figure 4A). VBNC had significant lower global metabolic activity as measured by the level of respiration determined from the metabolization of each compound (median= 124.0 [94.0-173] and 87.0 [65.0-118.8]) compared to 8D-NORMOx (246.0 [106.8-260.0] and 248.5 [123.0-262.0] for the two biological replicates) and to STAT (262.0 [136.5-272.0]) (p<0.01, Figure 4B). We identified two compounds (trimetaphosphate and adenosine-2’, 3’ cyclic monophosphate) that were more metabolized by VBNC than by control yeasts (Supplementary Table 1).

**Figure 4:**
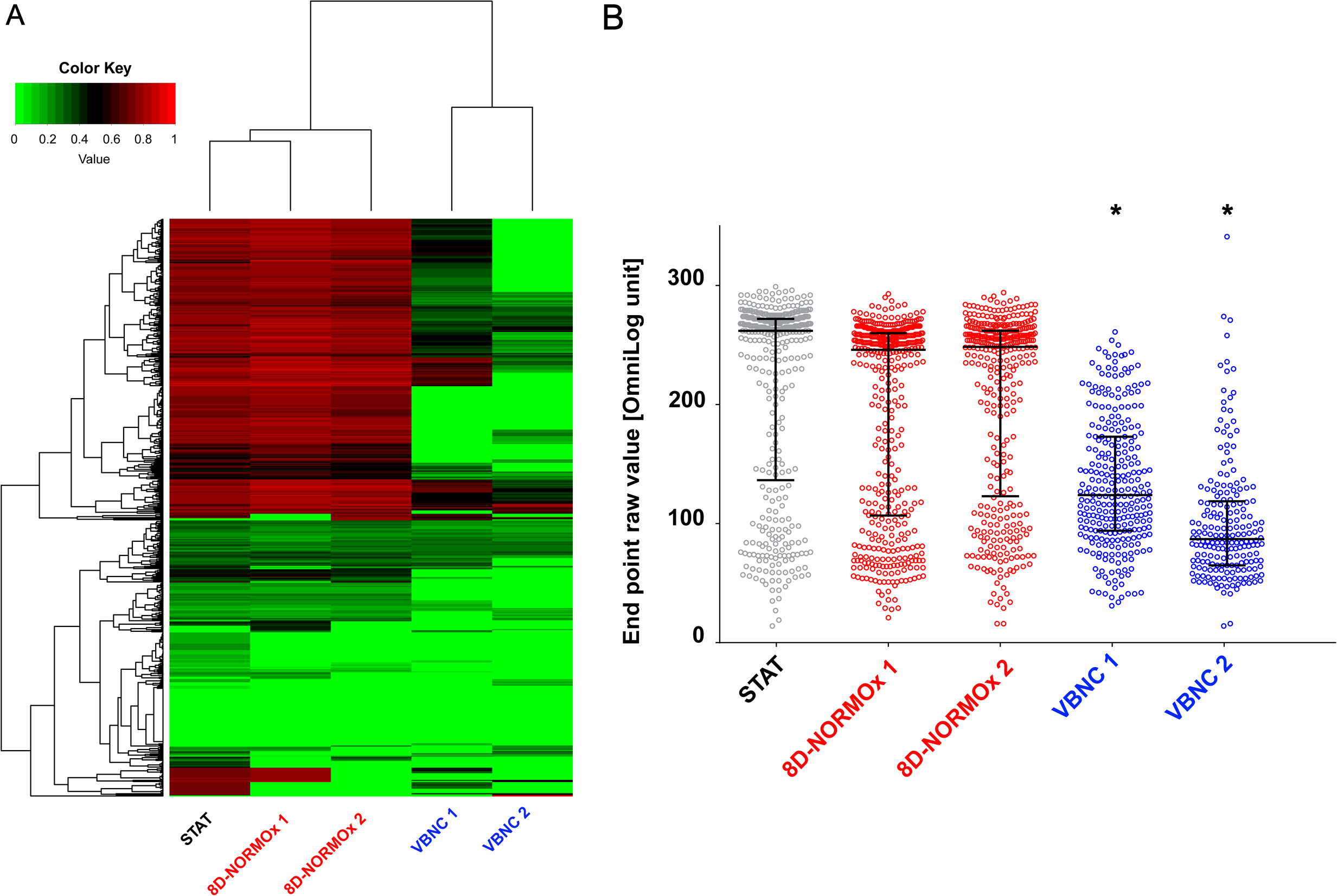
Phenotypic array analysis of VBNC showed a lower metabolic activity. We used the Biolog^®^ technology to explore the metabolic requirements of VBNC compared to STAT and 8D-NORMOx cells. We selected 8 plates and 761 different metabolites to scan the ability of substrates to be used as an energetic source. The respiratory activity (measurement of tetrazolium reduction in arbitrary units) was compared at the end of the incubation (6 days). Two biological replicates (1 and 2) were done for the VBNC/8D-NORMOx conditions. **A.** The heatmap of 521 selected compounds showed that VBNC exhibited a different metabolic profile than STAT and 8D-NORMOx. Color key corresponds to the normalized value of the OD between 0 and 1. **B.** The final endpoint analysis of the level of tetrazolium reduction (colorimetric change in tetrazolium dye or Omnilog unit, equivalent of the Optical Density measured at 590nm) revealed a lower activity for VBNC cells compared to cells in other conditions. VBNC and 8D-NORMOx conditions were done in duplicate (* p<0.01).

### VBNC have increased mitochondrial mass and depolarized mitochondria

Since the phenotypic microarrays based on mitochondrial activity measurement showed a decrease in VBNC, we further tested various mitochondrial parameters. Median ROS (152.3 [148.1-156.2] vs. 174.1 [168.9-177.8) and RNS (87.1 [78.2-101.2] vs. 149.5 [98.7-206.5]) productions were significantly decreased in VBNC compared to STAT, respectively (p<0.01, Supplementary Figure 5).

Using multispectral flow cytometry, we observed an increase in the mitochondrial mass in VBNC and in 8D-NORMOx compared to STAT and dead yeast cells (Mitotracker staining). Mitochondria in VBNC were mostly depolarized (low TMRE and high JC-1 stainings) (Figure 5A-B). Expression of the mitochondrial genes CYTb, NADH, mtLSU and COX1 significantly increased in VBNC compared to STAT (p=0.029) (Figure 5C), while DNA concentrations remained similar (p<0.05), suggesting that mitochondria are in high quantity, transcriptionally active, although depolarized.

**Figure 5:**
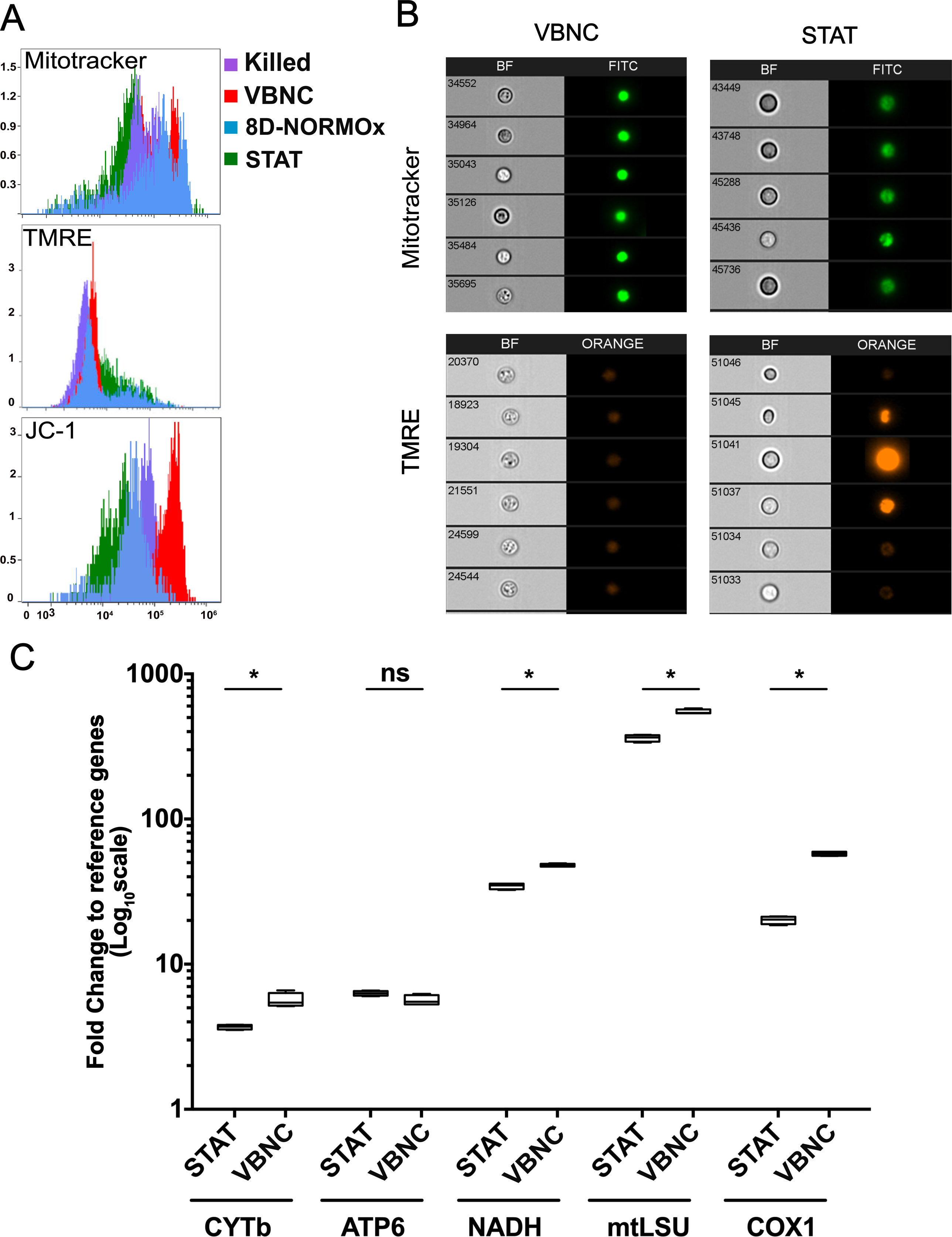
Mitochondria of VBNC were depolarized with an increased mass and gene expression. **A.** Mitochondria mass, and polarization were assessed in VBNC, 8D-NORMOx, STAT and heat killed yeasts using Mitotracker green and TMRE and JC-1 stainings, respectively and the ImageStream technology. In VBNC (red histogram), the Mitotracker staining is high, TMRE low and JC-1 high compared to STAT (blue histogram). Representative of two independent experiments. **B.** Microscopic visualization with Imagestream of the Mitotracker and TMRE stainings in VBNC and STAT yeasts, showing the contrast in the intensity of the stainings between conditions. Representative of two independent experiments. **C.** Expression of five mitochondrial genes, cytochrome B (CYTb), ATP synthase 6 (ATP6), NADH deshydrogenase (NADH), mitochondrial large sub unit (mtLSU), and cytochrome oxidase (COX1) as expressed as the fold change to the geometric mean of the expression of GAPDH and ACT1 housekeeping genes. The expression of CYTb, NADH, mtLSU and COX1 is significantly increased in VBNC compared to STAT (* p=0.029). Two technical and 2 biological replicates are represented here.

### VBNC have a reduced but a specific secretome and proteome

Proteins secreted in the supernatant (secretome) and cellular proteins (proteome) were compared in STAT, VBNC and 8D-NORMOx using both qualitative and quantitative analysis of the proteins recovered from experiments performed in biological triplicates.

We first focused on the differences between those conditions without considering the whole kinetics but only the initial point (STAT) and the final points (VBNC and 8D-NORMOx). A total of 1365 proteins were identified within the secretome and 3772 proteins within the proteome. In the secretome, the median number of proteins identified in STAT (747 proteins [598-797]) decreased over time in VBNC (411 [406-460] at D8) and not in 8D-NORMOx (910 [909-928]) (Supplementary Figure 6A, p <0.001). The protein concentrations in the secretome remained stable in VBNC and slightly increased in 8D-NORMOx (Supplementary Figure 6B, Supplementary Table 2). In the proteome, the median number of proteins identified in STAT (2975 proteins [2951-3029]) was stable over time, and evaluated at 3057 [3033-3171] in VBNC and 2968 [2960-2976] in 8D-NORMOx (Supplementary Figure 6C). The protein concentration in the proteome (0.005 mg/mL [0.002-0.011] in STAT) tended to increase in both hypoxia and normoxia reaching 0.017 mg/mL [0.008-0.018] and 0.021 mg/mL [0.016—0.027] in VBNC and 8D-NORMOx, respectively (Supplementary Figure 6D). We then analyzed the proteins that differed between STAT and VBNC or 8D-NORMOx in the secretome and the proteome. Overall, 17/1365 proteins present in the secretome, and 107/3772 present in the proteome were specific of VBNC (Supplementary Figure 7A-7B and Supplementary Table 3). When secreted proteins and cellular proteins were pooled, the analysis found 226 proteins present (Figure 6A). A total of 1654 proteins were found exclusively in the proteome common to the 3 conditions, or exclusively in STAT (n=150), in 8D-NORMOx/normoxia (n=75) or in VBNC/hypoxia (n=103) (Figure 6A). A total of 17 proteins were detected exclusively in the secretome common to the 3 conditions with 6 exclusively in STAT and VBNC (but different), and 16 exclusively in 8D-NORMOx (Figure 6A, Supplementary Table 4).

**Figure 6:**
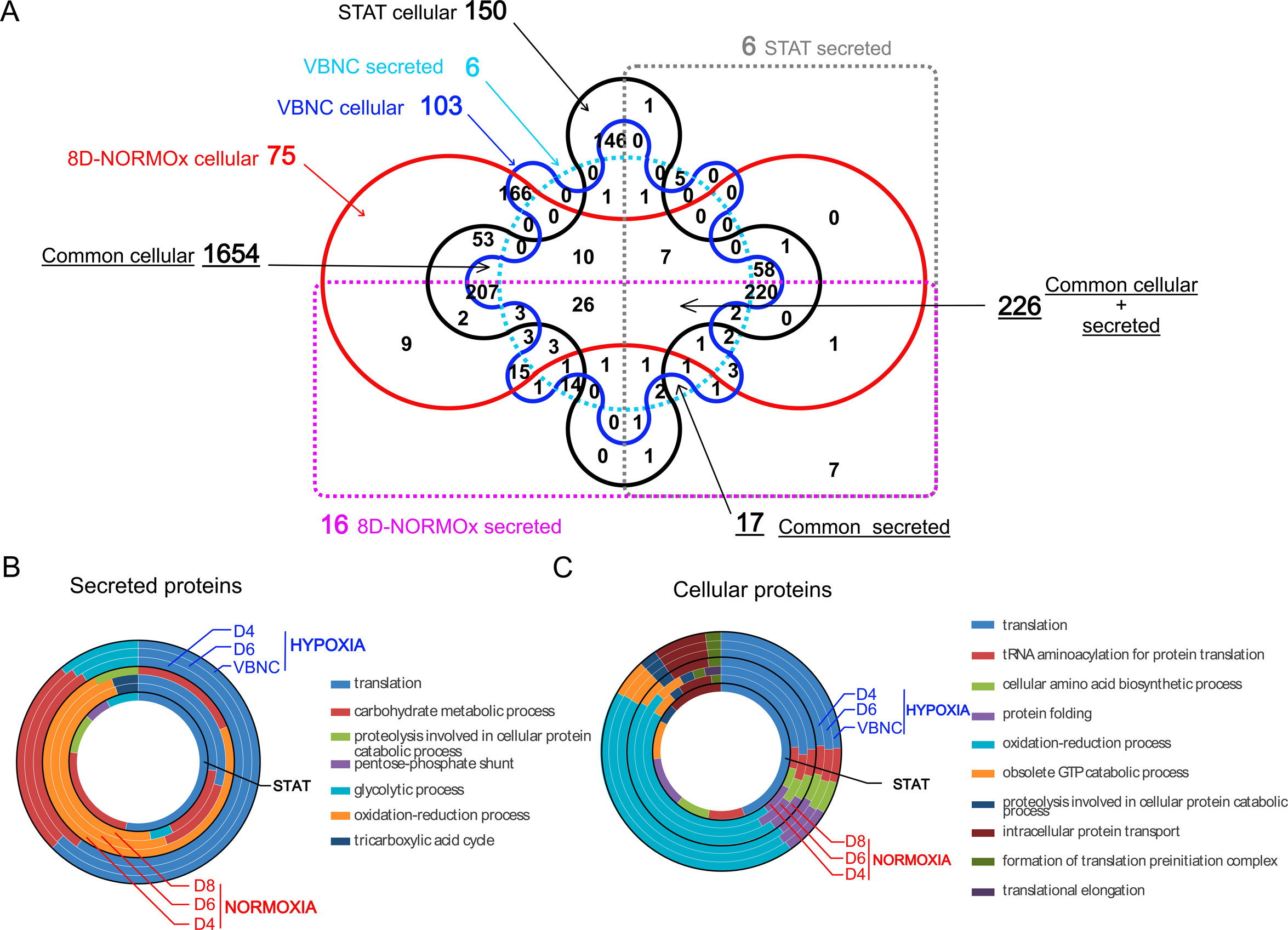
Secretome and proteome qualitative analysis showed specific features in VBNC. **A.** Venn diagram of cellular and secreted proteins in stationary phase (STAT), VBNC and 8D-NORMOx. Six secreted (clear dashed blue) and 103 cellular (deep blue) proteins were detected specifically in VBNC. Six were secreted only in STAT (grey dashed line) and 150 were detected in cellular protein only (dark line), 16 were secreted (pink dashed line) and 75 cellular (red line) only in 8D-NORMOx. Only proteins present in the 3 replicates for each condition were selected and compared. **B.** GO enrichment analysis of secreted proteins showed specific profile in VBNC. The major biological processes enriched in VBNC were translation, metabolic process of carbohydrates and glycolytic biological processes whereas oxidation-reduction was mainly enriched in 8D-NORMOx. **C.** GO enrichment analysis of cellular proteins showed similar profiles in VBNC and 8D-NORMOx with an enrichment for translation and oxidation-reduction.

For the subsequent analysis, the whole kinetics in hypoxia and normoxia was analyzed [STAT, hypoxia (D4, D6 and D8/VBNC) and normoxia (D4, D6 and D8/8D-NORMOx]. GO enrichment analysis of the proteins present in the secretome showed differences in the biological processes, molecular functions and molecular components, whereas similar patterns and distributions were observed for the proteome. Thus, the major enriched biological processes in the secretome were translation and metabolic processing of carbohydrates in hypoxia, while in normoxia they were related to oxidation-reduction mechanisms with no major change over time (Figure 6B). The enriched molecular functions and components were also different (supplementary Figure 8A and B and Supplementary Table 5). In the proteome, in both hypoxia and normoxia, the translation, oxidation-reduction biological processes (Figure 6C), the nucleotide binding transferase activity and the catalytic activity for molecular functions and the cytoplasm, ribosome, ribonucleoprotein and intracellular components were significantly enriched with no change over time (Supplementary Figure 8C and D and Supplementary Table 5).

**Figure 7:**
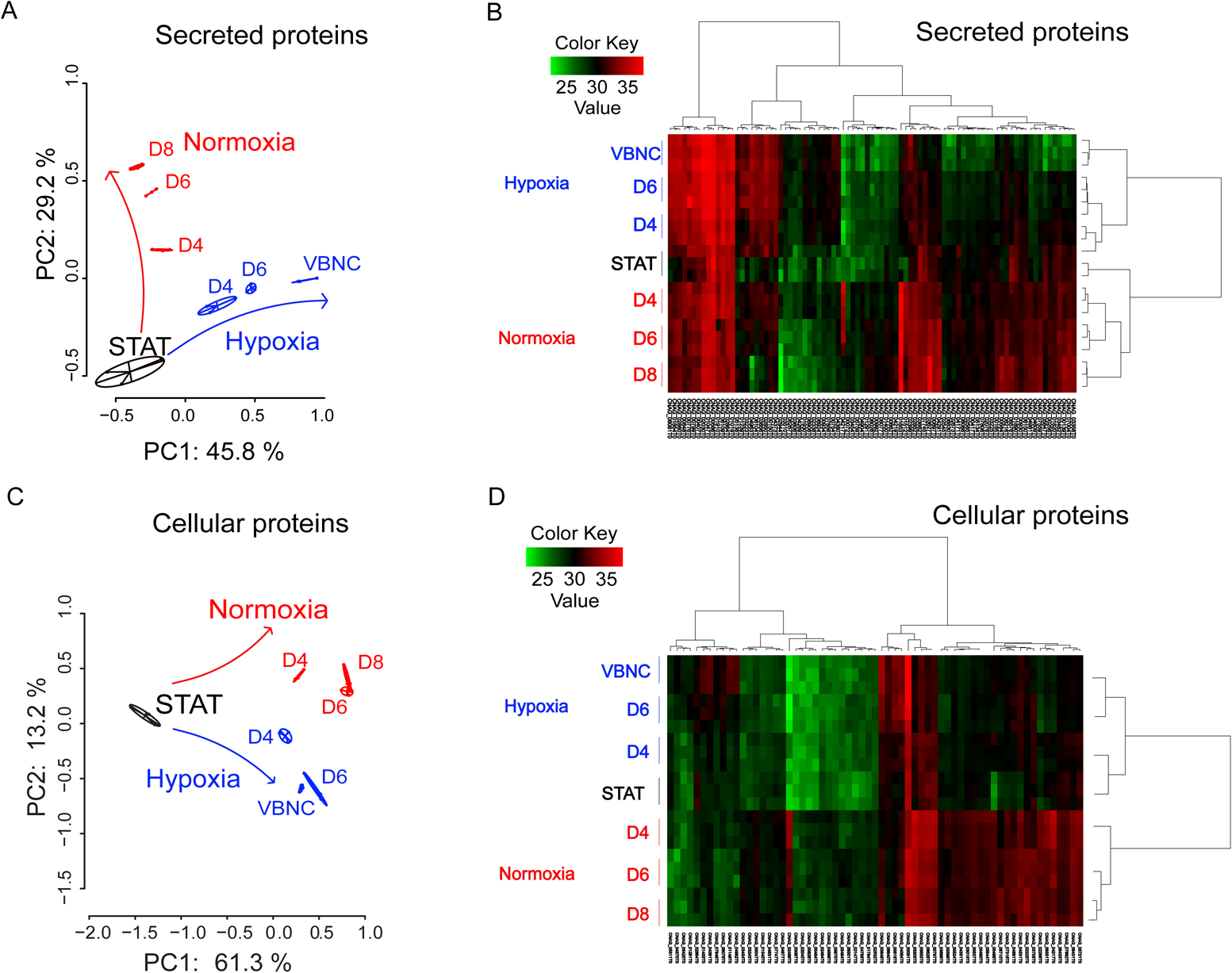
Secretome (A,B) and proteome (C,D) quantitative analysis showed an independent and specific kinetic evolution associated with VBNC generation. Principal coordinate analysis (PCoA) of secreted **A.** or cellular **C.** proteins and heatmap obtained by Limma analysis of secretome **B.** and proteome **D.** data. For the secretome, 85 proteins were differentially expressed overtime, including 49 significantly overexpressed in control and 36 in hypoxia. For the proteome, 63 cellular proteins were differentially expressed during kinetics including 47 significantly overexpressed in normoxia and 16 in hypoxia. Inversely of **B**, Data sets **D** showed that the STAT and hypoxia are closer than to normoxia. Biological triplicates were used for analysis.

**Figure 8:**
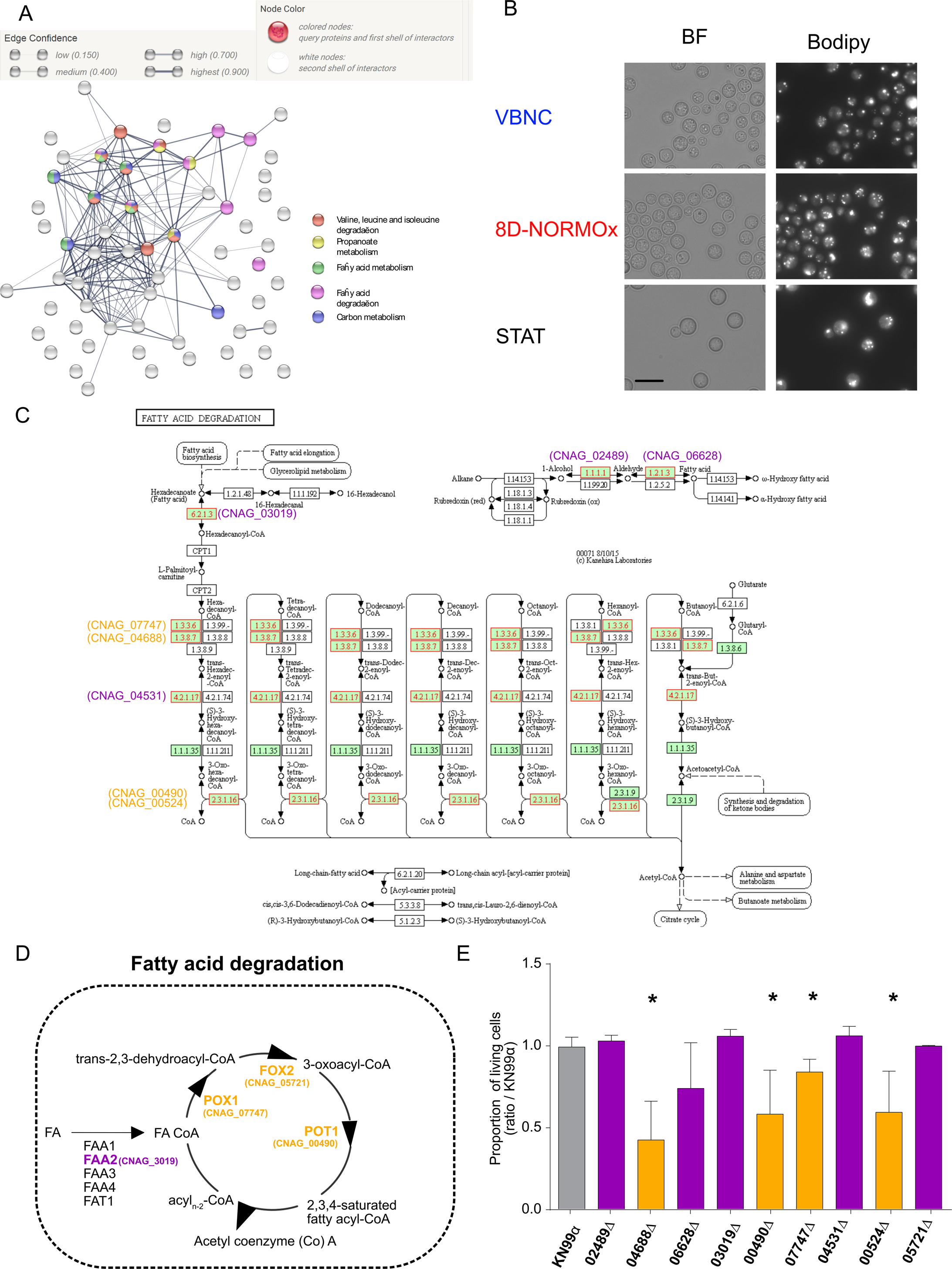
The fatty acid pathway participates in VBNC metabolism. **A.** The Search Tool for the Retrieval of Interacting Genes/Proteins (STRING) allowed the visualization of predicted protein-protein interaction for the 63 proteins hypoxia-regulated identified. It revealed an impact of hypoxia on diverse proteins as well as a prominent influence on fatty acids metabolism. Different clusters were identified from the network mapping: pathways linked to fatty acid metabolism (valine leucine and isoleucine degradation, propanoate metabolism, fatty acid degradation and metabolism) and to carbon metabolism. **B.** Lipid droplets stained with Bodipy^®^ 505/515 showed the presence of lipid droplets in all conditions in the same quantity. **C.** Eight entries among 63 proteins regulated in hypoxia mapped the fatty acid degradation by using KEGG pathway mapping (“map” pathways are not colored, *Cryptococcus*-specific pathways are colored green, the 8 entries are colored red) **D.** The fatty acid degradation pathway reconstructed from *Saccharomyces cerevisiae* pathway. FA (Fatty acid), FAA2 (Acyl Coa synthetase), POX1 (Fatty-acyl coenzyme A oxidase), FOX2 (3-hydroxyacyl-CoA dehydrogenase, 3-hydroxyacyl-CoA dehydrogenase), POT1 (3-oxoacyl CoA thiolase). **E.** Nine deletion mutants including those involved in the Acyl Coa synthase (purple bars) or in the fatty acid degradation (orange bars) pathway selected from KEGG database and the orthology with *S. cerevisae* were assessed for their ability to survive nutrient starvation and hypoxia in our conditions. The ratio of living cells for the mutants to those obtained with KN99α was determined based on LIVE/DEAD staining after VBNC generation (8D-HYPOx) as in Figure 2A. A significant decreased proportion of viable cells was measured for 04688Δ, 00490Δ, 07747Δ and 00524Δ (*p<0.01)

Principal coordinate analysis (PCA) of the global data revealed that the secretome (Figure 7A) and the proteome (Figure 7C) were modified in hypoxia and normoxia conditions compared to STAT. A specific pattern was seen as soon as D4 in hypoxia and normoxia. We selected the proteins present in all conditions and analyzed whether and how the level of each protein changed over time. Overall, 85 secreted proteins (including 49 in normoxia and 36 in hypoxia), and 63 cellular proteins (including 47 significantly more produced in normoxia and 16 in hypoxia) were differentially produced (Supplementary Table 6). Based on these results, hierarchical clustering of the secretome (Figure 7B) and the proteome (Figure 7D) was performed. STAT clustered together with the normoxia specimens for the secretome, whereas they clustered with the hypoxia specimens for the proteome.

Among the 16 cellular proteins overproduced during hypoxia, one was a serine carboxypeptidase (CNAG_06640); one was associated with intracellular transport (CNAG_07846); six were associated with the processing of DNA / RNA and the cell-cycle (CNAG_01455, CNAG_04570, CNAG_00819, CNAG_06079, CNAG_05311 and CNAG_04962), and the 9th protein was linked to the mTOR pathway (CNAG_01148, also known as FK506-binding protein) (Table 2). The other 7 proteins have no known roles (hypothetical proteins). Cellular proteins overproduced in normoxia were subsequently, down produced in hypoxia. These 47 proteins down produced in hypoxia were mostly proteins from the fatty acid degradation pathway, the glyoxylate cycle: isocitrate lyase (CNAG_05303), malate synthase (CNAG_05653) and a protein of the neoglucogenesis, the phophoenolpyruvate carboxykinase (CNAG_04217).

**Table 2:**
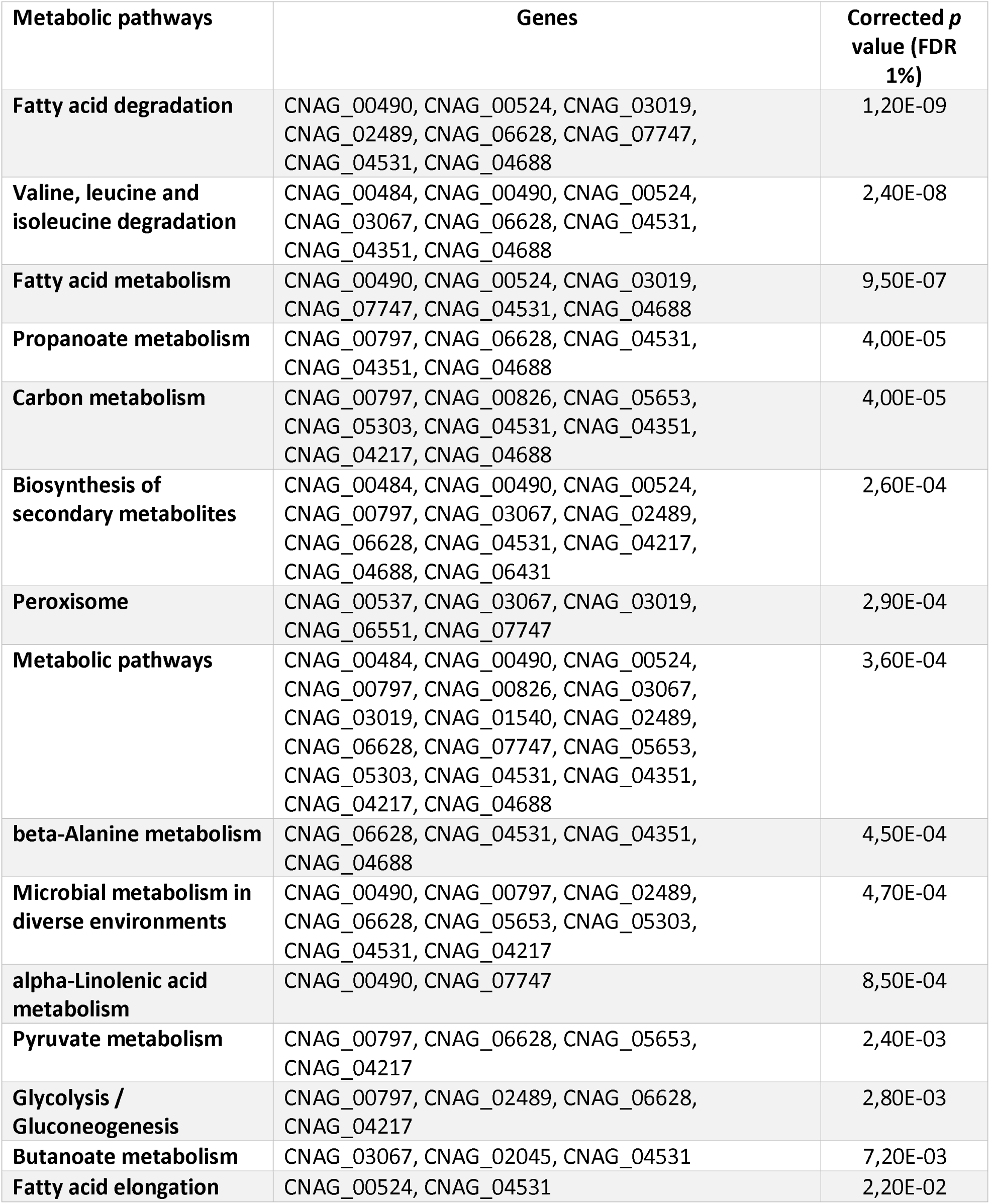
Metabolic pathways regulated in VBNC by STRING analysis

### The fatty acid pathway participates in VBNC metabolism

STRING analysis was used to visualize the predicted protein-protein interactions for the 63 cellular proteins differentially produced by VBNC (Figure 8A). Several metabolic pathways were identified, but especially the fatty acids metabolic pathways: degradation and metabolism of fatty acids, degradation of α-ketoacids (valine, leucine and iso-leucine), metabolism of alpha-linolenic acid, beta-alanine and elongation of fatty acids as well as peroxisomes. To examine whether VBNC stored fatty acids better than STAT or 8D-Normox, Bodipy^®^ 505/515 that stains neutral lipids in lipids droplets was used but no difference was observed (Figure 8B). Indeed, based on a manual count on a large number of cells (157 cells in STAT, 642 in VBNC and 392 in 8D-NORMOx), the mean dot/cell were respectively 3, 2.91 and 2.77, reaching no statistically significant difference. In addition, two independent flow cytometry experiment measuring the total excitation on 10 000 cells did not show differences in the geometric mean fluorescence intensity between STAT and VBNC. The β-oxidation pathway described in the KEGG database was used to visualize the 63 cellular proteins differentially produced in VBNC and the 8 proteins for fatty acid degradation metabolism (Figure 8C). The fatty acid degradation pathway (β oxidation) is well-characterized in *Saccharomyces cerevisiae* with several genes found also in *C. neoformans* genome (Figure 8D). Interestingly, the corresponding proteins in *C. neoformans* were known to be under produced in VBNC. The 9 deletion mutants of proteins identified from the KEGG database and by the orthology with *Saccharomyces* genome available in the Madhani collection were tested in the model for their ability to generate VBNC. A significant decreased in the proportion of living cell was measured for 04688Δ, 00490Δ, 07747Δ and 00524Δ, all directly involved in fatty acid degradation (Figure 8E, p<0.01), suggesting that the fatty acid pathway is required in VBNC although the level of expression is lower than in STAT cells.

### VBNC cells have reduced but specific transcription

We then compared genes expression in the three conditions using transcriptome analysis. To avoid bias related to decreased RNA content and not in gene expression, we performed a specific experiment in which we spiked each sample with the same quantity of *S. cerevisiae* prior to extraction allowing us to normalize the transcription of *C. neoformans* to that of *S. cerevisiae*. The number of transcripts decreased in VBNC, and to a lesser extent in 8D-NORMOx, compared to STAT and LOG conditions that harbored similar level of transcripts (Figure 9A).

PCA analysis of the global transcriptomic data from non-spiked samples suggested an evolution from LOG to STAT phases with a clear independent and specific route leading to VBNC or 8D-NORMOx (Figure 9B). GO enrichment analysis of upregulated transcripts showed that the major biological processes involved were: signal transduction, ATP synthesis coupled proton transport and regulation of ARF protein signal transduction in VBNC, oxidation-reduction, protein folding and response to stress in 8D-NORMOx and oxidation-reduction and translation in LOG (Figure 9C). Further analysis revealed 7 clusters that clearly separate the transcriptional profile of the 4 experimental conditions (LOG, STAT, VBNC, 8D-NORMOx, Figure 9D). Specifically, cluster 4 appeared to be composed of genes upregulated in VBNC compared to the other conditions (underlying pathways summarized in Table 3 and Supplementary Table 7).

**Table 3:**
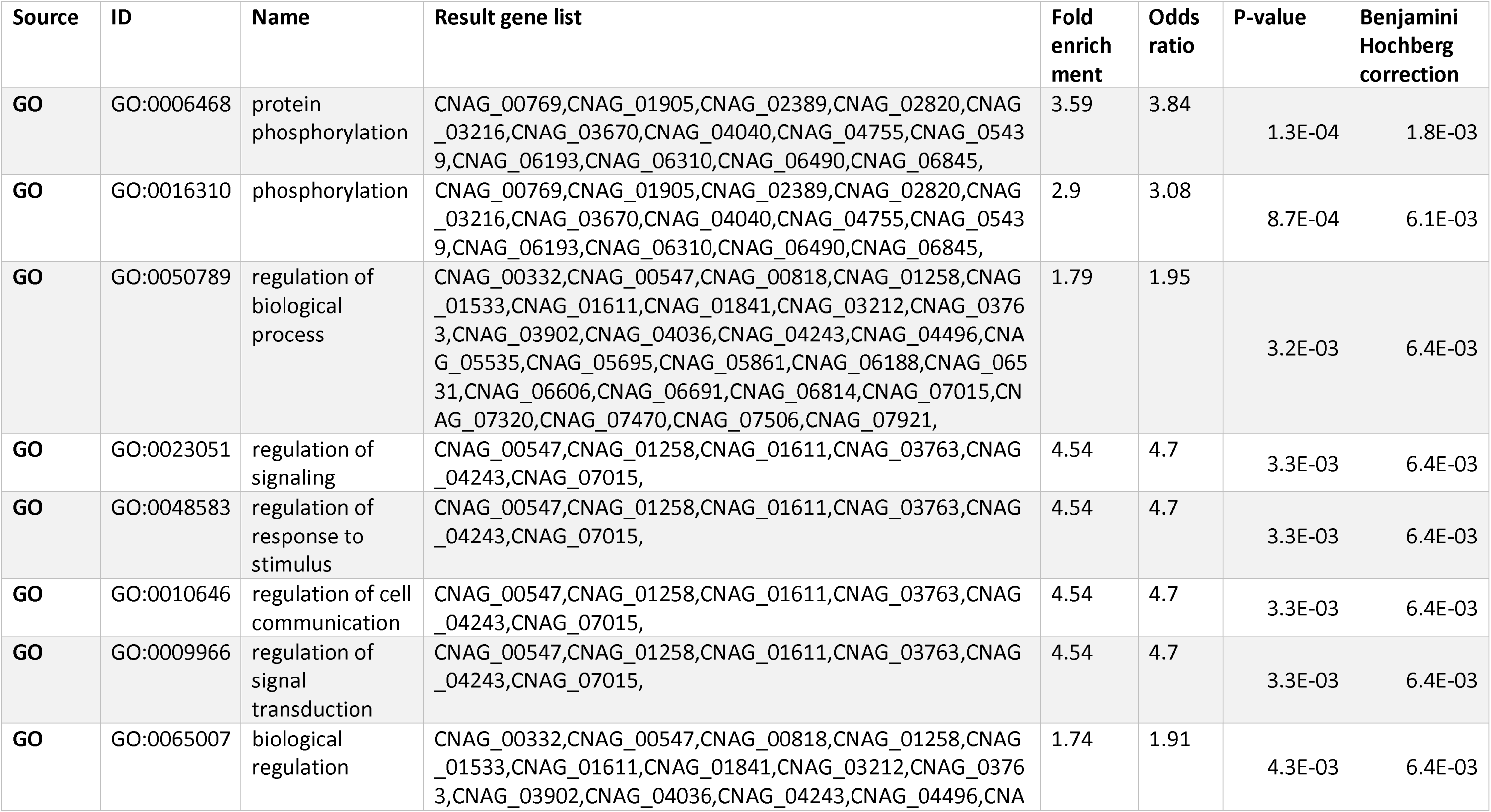
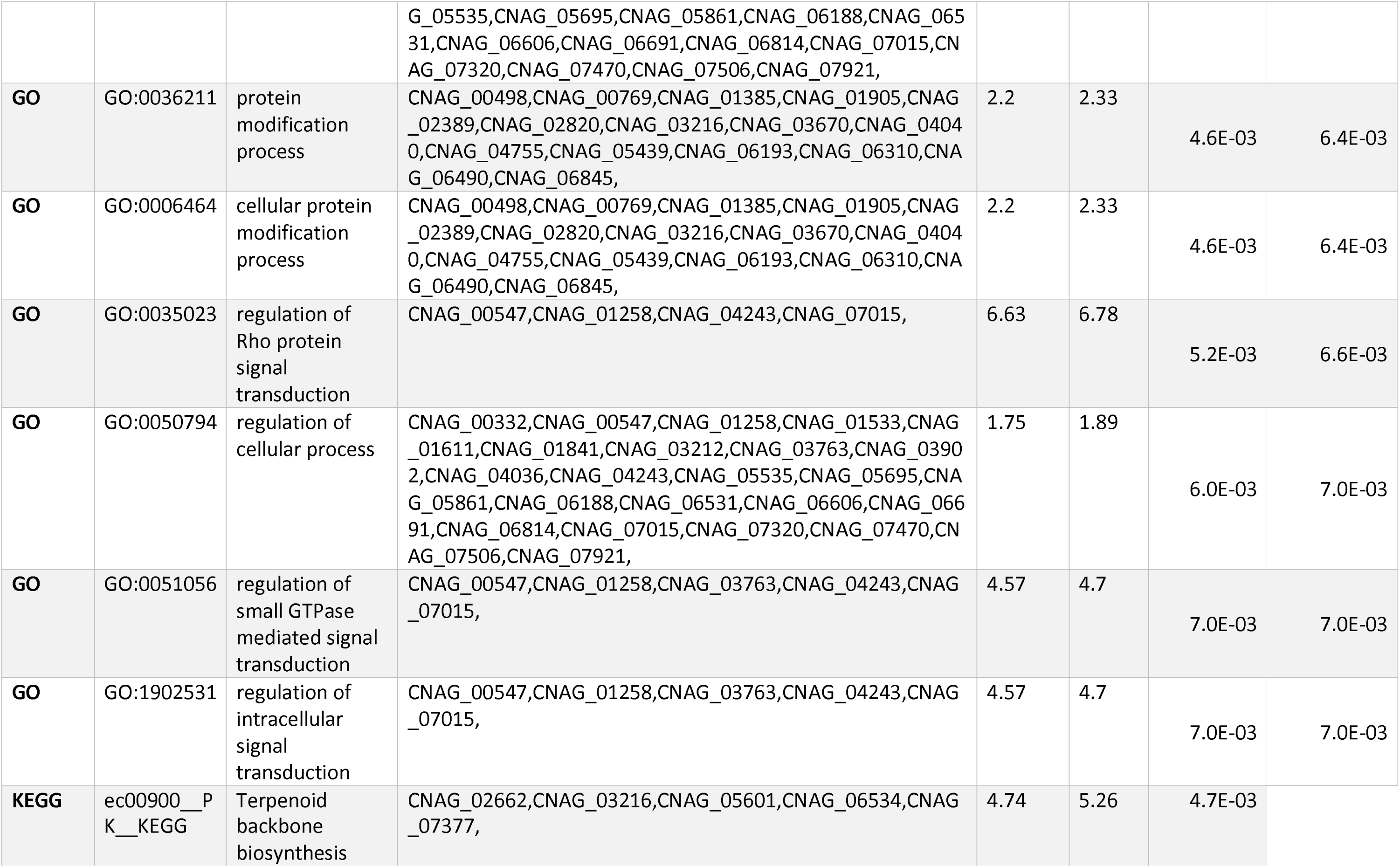
Transcriptomic pathways involved in the VBNC cluster 3 and genes the corresponding upregulated genes.

**Figure 9:**
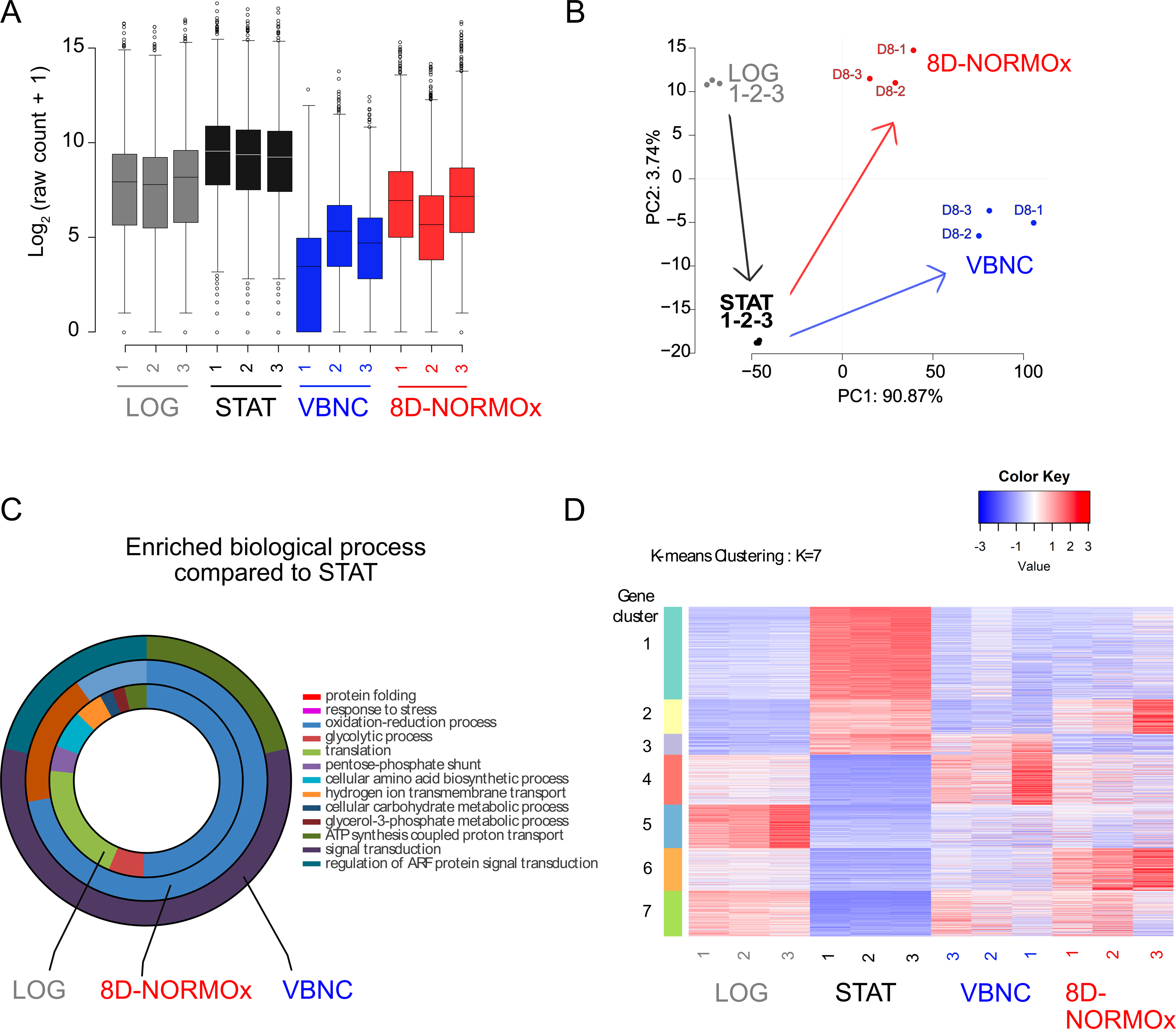
The global transcription is decreased in VBNC but remains specific. **A.** Transcriptional level of all conditions relative to that of *S. cerevisiae* spiked in each sample showed a significant decrease in VBNC compared to other conditions **B.** PCA analysis of the global transcriptomic data (wo_spikes) suggested that logarithmic cells (LOG) evolve to STAT which evolve to VBNC or 8D-NORMOx **C.** GO enrichment analysis of upregulated transcript (wo_spike) compared to STAT. The major enriched biological processes were signal transduction, ATP synthesis coupled proton transport in VBNC and protein folding, oxidation reduction process and response to stress in 8D-NORMOx. D. Heat map of the gene expression revealed the presence of 7 clusters displaying different expression patterns for all conditions. The color key represents normalized transformed counts. Biological triplicates were used for analysis.

## Discussion

We report the existence of a *C. neoformans* phenotype whereby dormant cells *in vitro* are viable but not culturable cells (VNBC). Hence, *C. neoformans* joins the ranks of microbes where similar phenomena have been described [6,8,10,12–16] with the proviso that for this organism this phenotype has obvious clinical importance given that latency is a common manifestation of human infection.

Latency of cryptococcosis that can be linked with dormancy of *C. neoformans* had been characterized clinically [30,31] and epidemiologically [17,18]. The dormant yeasts are thought to be located in a pulmonary granuloma but other reservoirs could exist [23,32]. Recently, evidence for the existence of dormancy in a subpopulation of *C. neoformans* upon interaction with the host *in vivo* (murine infection) and with macrophages *in vitro* have been provided [23]. To obtain an homogenous population of dormant cells in amounts allowing in depth characterization, we set up an *in vitro* protocol using stresses known to generate quiescence in other cell types, nutrient starvation and hypoxia [33–35]. Indeed, dormant yeasts recovered from murine infection are too scarce (<30/lungs) to work with and mixed with other cells requiring sorting that could impact their phenotype [23].

The standardized protocol generated yeasts that exhibited high viability (>99%), low apoptosis (<1%) [36] and low culturability (<0.01%) at D8 which are characteristics of VBNC. This phenotype is associated with dormancy in other microbial species/populations [6–8,10,37]. Hypoxia turned almost the entire *C. neoformans* population into a VBNC phenotype (<15% culturability at D2 suggesting that hypoxia triggers the switch towards VBNC rapidly. Hypoxia is an important factor inducing dormancy in mammalian stem cells [24]. In stem cells, hypoxia is required for a strict metabolic regulation to maintain long-term quiescence and self-renewal of populations. Hypoxia is also present to some extend in the granuloma [38–40]. Indeed, hypoxia is also known to influence cell fate (differentiation, de-differentiation) within the stem cell niche [41,42]. In addition, resistance to apoptosis is known in mammalian cells by the induction of apoptosis inhibitors such as IAP-2 [43,44] but opposite results have been found with tumor cells [45]. Nevertheless, by using these *in vitro* conditions to generate what happened to be VBNC, we acknowledge/are aware that this phenotype could be different from the subpopulation of dormant yeasts identified from the lung of infected mice [23], although similarities were observed (low stress response through CMFDA decreased fluorescence, latency of growth, increased mitochondrial transcription). One of the major characteristics of cells manifesting the VBNC phenotype is their ability to be reactivated - to divide again - in response to specific stimuli [7,46,47]. Given that the critical issue in reactivation experiments is to distinguish between more cells being able to grow and faster growth of the culturable cells already present, a method was developed to measure the probability of growth for one yeast cell, inspired by the most probable number method [48]. The determinant experiment to assess the VBNC phenotype was the demonstration that PA addition allowed reactivation (1.2‰ reactivated VBNCs). The effect of PA resulted from both the doubling number of reactivated VBNC and an increase in growth rate, without a clue on which factor was more important. It is highly probable though that resuscitation of VBNC requires other factors that need to be discovered. It is also possible that the majority of the VBNC may not be able to reactivate in the conditions used here as hypoxia may have pushed the yeasts too far into the process of dormancy [49], with a majority of cells injured while still alive but unable to divide again [50].

We additionally observed that MM dilutions increased growth capacities of VBNC (decreased latency. It did not influence VBNC reactivation but rather decreased the growth rate by a faster rewire of the metabolic state of the remaining culturable cells. To explain this counterintuitive observation, we propose that exposure to a high amount of nutrients after long-term starvation was deleterious for VBNC that had to resume normal metabolism, possibly as a result of oxidative damage as metabolism increases. This hypothesis is further based on the fact that death was shown by others to be accelerated in rich substrate [51–53].

Despite the importance of *Cryptococcus*-protozoan interaction in the explanation of the yeast’s virulence and intracellular pathogenic strategy [54], we found that macrophage uptake of VBNC had no effect on culturability. This result is different from what was observed for other microorganisms. For example, reactivation of VBNC was possible for the bacterium *L. monocytogenes* and the protozoans *Acanthamoeaba polyphaga* [55] and *A. castellanii* [56].

We first screened the dormancy phenotype through the decrease in metabolic and stress responses [23], as measured upon staining with the CMFDA fluorescent probe [26,57]. Although it allowed us to suspect that hypoxia was an important factor for the generation of VBNC, the equivalent culturability of subpopulations sorted by their fluorescence intensity suggested that CMFDA staining is not a perfect marker of metabolic and stress response in *C. neoformans*.

Decreased metabolic activity in VBNC was demonstrated by several strategies. First, a phenotypic array measurement (Biolog^®^) allowed us to show a reduction in the metabolism of 761 compounds in VBNC. Second, the concentration of secreted or intracellular proteins was lower in VBNC compared to 8D-NORMOx as reported during quiescence in other organisms [58–61]. In parallel, the increased number of proteins present in the supernatant of 8D-NORMOx could correspond to increased secretion but more likely to an increased release of proteins due to a high cellular mortality observed in that condition. Third, using a unique strategy of analysis for the transcriptome including the spike of a definite percentage of Log phase *S. cerevisiae* cells, we demonstrated a global decreased transcriptional activity in VBNC compared to 8D-NORMOx, LOG and STAT conditions, identifying again the lower metabolic activity of VBNC.

In VBNC, we found that the mitochondrial phenotype included an increased mitochondrial mass, increased depolarization and an increased expression of mitochondrial genes. This observation could explain why ROS and RNS levels were lower in VBNC compared to STAT. This depolarized state could also reflect activation of autophagy [62]. These results are reminiscent of those obtained in the subpopulation of dormant yeasts observed in mice 7 days after infection and in macrophages, where *COX1* and autophagy genes (*ATG9* and *VPS13*) are upregulated compared to other subpopulations of yeasts [23].

We identified and quantified proteins from the secretome (supernatant) and proteome (cell pellet) over time. Overall, 1365 secreted proteins were identified. This number is greater than previously described [63–65] but the protein extraction methods differed (gel extraction, TCA / acetone precipitation alone), and the identification procedures as well as the culture media used (YPD or MM) were different. Nevertheless, of the 191 secreted proteins previously identified in stationary phase in YPD [63–65], 93.7% were found in our corresponding sample (STAT with 565 proteins identified in all of the 3 replicates). In *C. neoformans*, the secreted molecules can take several pathways to reach the periphery of the cell and the extracellular space. There are conventional and unconventional secretory pathways involving secretory machinery in protein export: Sec4, Sec6, Sec14, the Golgi apparatus, and extracellular exosome vesicles [66–68]. The TCA precipitation step used here to analyze the secreted proteins within the supernatants precipitates soluble proteins and also those in extracellular vesicles [69]. Consequently, we are unable to precisely identify their mechanism of secretion. This opens future directions for studying the relationship between dormancy and extracellular vesicles. A secreted protein, pqp1 peptidase (CNAG_00150) was found overexpressed in hypoxia. This peptidase cleaved the pro-qsp1 quorum sensing peptide into qsp1 (CNAG_03012), that acts on the whole cell population and regulates virulence [70–72]. However, we did not find qsp1 in the supernatants of the culture media and yet it is present in micromolar amounts in the publication of Homer et al. [70]. It is possible that our extraction protocol was not optimized for small peptides (11 amino acids) or that small peptides might have been eliminated through the various washing steps. However, with our optimized extraction method, we identified more cellular proteins (n=3772) than previously reported in *C. neoformans* [64,73–75]. From the quantitative proteome analysis of VBNC, we identified the FK506-binding protein which is involved in the mTOR pathway, and known to play a role in quiescence in *S. cerevisiae* [76]. This protein is able to bind to rapamycin (Sirolimus^®^) in both yeasts and mammalian cells promoting quiescence [77,78]. In *C. neoformans*, rapamycin induces cell cycle arrest in the G1 phase [79].

The STRING analysis of the protein-protein interactions provided evidence that the regulation of fatty acids was the major metabolic pathway involved in VBNC. The regulation of fatty acids is a phenomenon that occurs in *S. cerevisiae* quiescence by participating in energy cellular homeostasis and membrane synthesis [33]. In *Vibrio vulnificus*, changes in fatty acid membrane composition contribute to maintaining the viability of the VBNC [37]. A common feature among the metabolic fatty acids pathways is the participation of acetyl-CoA [80]. Acetyl-CoA is more than a cofactor in cell metabolism. It is central to the physiology of mammalian cells (fatty acid metabolism, Krebs cycle, apoptosis, cell cycle, damage-response DNA and epigenetics) [81]. In *C. neoformans*, the genes of the three principal production routes of acetyl-CoA in the cytosol are regulated [82]: the β-oxidation of fatty acids by the enzyme Mfe2, acetate by acetate synthase Acs1 and citrate by Acl1. Our hypothesis is that maintenance of fatty acid metabolism in hypoxia would allow the cells to maintain membrane integrity and viability, with an increased mortality in deletion mutants in key enzymes of the fatty acid pathway. If these results are in favor of the indispensability of acetyl-CoA for the maintenance of *C. neoformans* infection, the relationship between acetyl-CoA, PA – a precursor of acetyl-CoA – and dormant cells should be further analyzed. The fact that PA was able to resuscitate VBNC cells, confirms the potential relevance of the link made by the proteomic study between VBNC and metabolic pathways of acetyl-CoA. Qualitative and quantitative changes in fatty composition occurs in *C. neoformans* depending on the growth phase, with an increase during growth progression [83]. In addition, KEGG pathway mapping of the 63 proteins regulated in hypoxia showed an involvement of 8 proteins involved in fatty acid degradation. The 8 proteins involved were downregulated in hypoxia compared to normoxia. Another reconstruction of the fatty acid degradation pathway using *Saccharomyces* genome database involved the 4 proteins also downregulated in hypoxia *FAA2* (CNAG_03019), *POX1* (CNAG_07747), *FOX2* (CNAG_05721), *POT1* (CNAG_00490). The mutants *pox1Δ, pot1Δ, CNAG_04688Δ* and *CNAG_00524Δ* tested here had a decreased viability in hypoxia compared to the parental strain KN99α strengthening the role of fatty acid degradation in hypoxia and emphasizing the importance of downregulating fatty acid degradation in VBNC phenotype. In *Candida albicans* macrophages internalization also results in the induction of fatty acid degradation pathways [84] and similar patterns of expression are observed in *C. neoformans* in the lungs of infected mice [85] and in response to stress within macrophages and amoeba [86].

In summary, we describe conditions that induce *C. neoformans* to switch to a homogenous population of viable but non culturable cells that can be considered dormant and use those conditions to study the biological characteristics of these yeasts by different approaches. VBNC cells manifest what is potentially an extreme phenotype (dormancy) derived from stationary phase (quiescence) but demonstrate the production of a very specific secretion pattern critical to the pathobiology observed in the mammalian hosts. Our study identified genes involved in the emergence and maintenance of the VBNC phenotype, which implicate a major role for lipid metabolism. The availability of conditions that reliably induce VBNC provides a new research tool in the field of fungal and specifically *C. neoformans* biology to study the important features of dormancy and help understand latency in cryptococcal pathogenesis.

## Material and methods

### Culture media and strains

*Cryptococcus neoformans* strains (Supplementary Table 8) were usually grown in liquid Yeast Peptone Dextrose (YPD: 1% yeast extract (BD Difco, Le Pont de Claix, France), 2% peptone (BD Difco), 2% D-glucose (Sigma, Saint Louis, Minnesota, USA)). Minimal medium (MM, 15mM D-glucose (Sigma), 10 mM MgSO_4_ (Sigma), 29.4mM KH_2_PO_4_ (Sigma), 13mM Glycine (Sigma), 3.0 µM Thiamine (Sigma), pH5.5) was used for specific experiments [29].

### In vitro generation of C. neoformans VBNC

*Cryptococcus neoformans* H99O strain stored in 20% glycerol at −80°C was cultured on Sabouraud agar plate at room temperature (step 1). After 2 to 5 days of culture on agar plate, 10^7^ yeasts (obtained with a calibrated 1µL loop) were suspended in 10 mL of YPD in a T25cm^3^ flask with vented cap disposed up in the incubator and cultured for 22 hours at 30 °C, 150 r/min with lateral shaking until stationary phase (final concentration≈2×10^8^/mL) (step 2). One hundred μL of this first culture was added to 10 mL YPD (second culture) and incubated in the same conditions until stationary phase (Step 3). In stationary phase, the medium is deprived of nutrient (carbon and nitrogen) thus preventing yeast growth. Then, the flasks were placed in a GENbag^®^ (Biomérieux) anaerobic atmosphere generator to generate hypoxia (< 0.1% O_2_) and incubated for 8 days in the dark at 30°C. These yeast cells are called 8D-HYPOx and then VBNC for convenience. Control samples consisted of yeasts in stationary phase (Step3, STAT) and yeasts incubated under the same conditions but in normoxia are called 8D-NORMOx (Step 4) (supplemental Figure 1). Of note, the atmosphere in the bag is enriched in CO_2_ by the biochemical process of oxygen depletion.

### Transmission electron microscopy

Freshly harvested cells from STAT and 8D-HYPOx were fixed overnight at 4°C with 2.5% glutaraldehyde in 0.1 M sodium cacodylate buffer (CB) at pH 7.2. Then samples were washed in 0.1 M CB (pH 7.2) and post-fixed in a 1% osmium tetroxide in CB at room temperature for 1 h. We pre-embedded the cells in 4% agar type 9: the cells were mixed with agar, spun-down by gentle centrifugation and the pellet stored at 4°C until the agar had solidified. We excised 2 mm^3^ sections from agar block, and processed them in the automatic MicroWave tissue processor for electron microscopy (AMW, Leica Microsystems, Vienna, Austria) as recommended by the manufacturer: samples were washed with water and dehydrated with graded ethanol concentrations (25–100%), followed by a mixture of graded resin Spurr Agar low viscosity Resin (Agar scientific, Gometz la ville, France) concentration in ethanol (35-100%). The embedded agar blocks were polymerize at 60°C for 48h. Ultrathin sections (70 nm) were performed with an ultramicrotome ‘Ultracut UC7’ (Leica Microsystems, Vienna, Austria), stained with 4% uranyl acetate and Reynold’s lead citrate onto 100 mesh copper grids coated with a thin glow carbon film (S162-1, Oxford Scientific, Eynsham, UK) and then observed under Tecnai SPIRIT (FEI-Thermofisher Company) at 120 kV accelerating voltage on a camera EAGLE 4Kx4K (FEI-Thermofisher Company).

### Stainings and dyes

Unless otherwise stated, yeasts were washed in phosphate buffered saline (PBS) twice before and 3 times after staining. Stress response was assessed by measuring glutathione levels using 5-chloromethylfluorescein diacetate (Cell tracker^™^ green CMFDA, Life Technologies) as already shown in mammalian cells [26,87] and yeasts [23,27]. Yeasts (10^6^) were labeled with 3.3μM CMFDA in 500μL of PBS for 30 min at 37°C in the dark. Anti-glucuronoxylomannan antibody binding on the capsule was performed with E1 monoclonal antibody, as described previously [88]. Nuclei were stained using 41,6-diamidino-2-phenylindole (DAPI, Invitrogen, Carlsbad). Yeasts (10^7^) were fixed with 3.7% paraformaldehyde for 30 min before staining with 0.5µg/mL of DAPI for 5 m. MDY-64 (Thermofischer Scientific, Waltham) was used to stain the intracellular vacuolar membranes by incubating 10^7^ yeasts for 5 min with 5µM MDY-64. Lipid droplets were stained using Bodipy^®^ 505/515 (Invitrogen, Carlsbad) as already used in *Saccharomyces cerevisiae* [89] at 1µg/mL with 10^7^ cells for 20 min at room temperature. To assess viability, LIVE/DEAD^®^ Fixable Violet Blue (416 nm/451 nm) diluted at 1:1000 and 250 µL was added to 10^7^ yeasts pellets and incubated at 30°C in the dark. Controls were dead yeasts generated upon exposure to 200 mM of oxygen peroxide (H_2_O_2_) at 37°C, under 650 r/min agitation or heat killed 1h at 65°C.

Yeasts were analyzed by fluorescence microscopy and pictures taken using an AxioCam MRm camera (Carl Zeiss, Oberkochen) on interferential contrast microscope (DMLB2 microscope; Leica, Oberkochen) with Zeiss Axiovision software.

Mitochondrial mass and polarization were evaluated with Mitotracker Green (Invitrogen) at 40 nM, JC-1 (Invitrogen) at 10 µg/ml and TMRE at 200nM in PBS on 10^7^ yeasts for 30 min at 37°C and then washes 3 times in PBS. Reactive oxygen (ROS) nitrogen species (RNS) measurements were also performed. Microplate wells were filled with 10^4^ yeasts in 100 μL PBS. Briefly, the probes for ROS (2’,7’-dichlorofluorescein diacetate, Sigma) and RNS (dihydrorhodamine 123, Invitrogen) were diluted at 100 μM in PBS and methanol, respectively, and 20 μL were added to wells of a microtiter plate containing 10^4^ yeasts in 100 μL of PBS. Hydrogen peroxide (4M) was included as a positive control. The plates were incubated at 37°C in the dark. After 1 hour, fluorescence was measured with a fluorometer (Fluoroskan ascent™ fluorometer, Thermos Fischer Scientific, Waltham, MA, USA) using excitation 485 nm and emission wavelengths of 530 nm [90]. The data were expressed as arbitrary units of fluorescence ±SD.

### Apoptosis assay

Yeasts (10^7^) were fixed for 1 h at room temperature in 3.2% formaldehyde (Sigma) in PBS. After washings in PBS, yeasts were incubated in 1 mL of 5% 2-mercaptoethanol in SPM buffer (Sorbitol 1.2 M (Sigma), potassium phosphate monobasic 50 mM (Sigma), magnesium chloride hexahydrate 1 mM (Sigma), pH 7.3) for 1 h at 37°C. Yeasts were washed once in SPM buffer and incubated for 1 h at 30°C in 1 mL of a solution containing 2 mg of Zymolyase 20T (Euromedex^®^), 100 mg of lysing enzymes from *Trichoderma harzianum* (Sigma) and 1 mg of BSA (Sigma) in spheroplasting buffer (sorbitol 1 M (Sigma), sodium citrate tribasic dihydrate 100 mM (Fluka), EDTA 10 mM (Sigma), pH 5.8). All these incubations were performed in the dark with agitation at 650 r/min in a thermomixer. The spheroplasts were then washed 3 times in SPM buffer permeabilized by incubation in 1 mL of 0.1% triton X-100 (Sigma) in PBS for 2 minutes on ice and washed twice in PBS. TUNEL-Fluorescein (In Situ Cell Death Detection Kit, Roche Life Science) was used to measure apoptosis at the single cell level following manufacturer’s recommendations. Yeasts were resuspended in 50 µL of solution of 10% of TdT and 90% dye-coupled dUTP in PBS, incubated 1 h at 37°C and then washed twice in PBS. Apoptosis control was generated by incubation of 3×10^7^ yeasts in 2 mM H_2_O_2_ for 3 h at 37°C. TdT was omitted for the negative control. In each run, yeasts treated with DNAse (3 U/mL, Ambion^®^) were used as a TUNEL positive control.

### Flow cytometry measurements and cell sorting

Flow cytometry was used to quantify MDY64 staining, viability and apoptosis assays, yeasts - using the Guava easyCyte 12HT Benchtop Flow Cytometer^®^ (Guava, MERCK, Kenilworth, New Jersey).

Multispectral flow cytometry (ImageStreamX, Amnis Corporation) was used to quantify mitochondrial fluorescence as described before [23,88]. For each population of interest, the geometric mean was calculated using the IDEAS software.

In specific experiments, yeasts were sorted on a BD FACS aria™ II flow cytometer. The gating strategy included the exclusion of the aggregates and the doublets. Subsequently, the sorting was carried out according to the intensity of CMFDA in 3 populations (CMFDA^high^, CMFDA^medium^ and CMFDA^low^).

### Growth curves method and statistical modelling

Growth curves measurements were done using the Bioscreen^®^ apparatus (Oy Growth Curves Ab Ltd). Yeasts suspensions (300 μL at 10^4^/mL) were added to wells of a plate and incubated for 2 to 5 days at 30°C with continuous, high amplitude and fast speed agitation. Optical densities (OD) were recorded at 600nm every 20 minutes. The R software with the Grofit plug-in [91] was used and for each replicate and in each condition, we adapted the best mathematical model according to the AIC (Akaike Information Criterion). The two models that best modeled the growth curves were the logistic and Richards modified, where λ represents the length of lag phase or latency of growth, *μ* the growth rate, A the maximum OD (concentration), *v* the shape parameter and t the time of incubation.

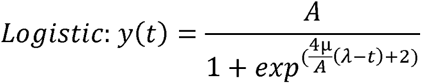

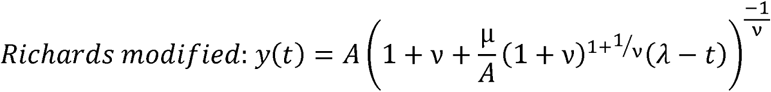

These formulas were written in Graphpad Prism v6.02 to allow curve extrapolation and determination of the latency parameter λ.

### Culturability by CFU method and plating method

To assess basic culturability, the suspensions recovered at the end of the assay were enumerated and 3000 cells, counted with the Guava easyCyte 12HT Benchtop Flow Cytometer^®^, were plated in duplicates on YPD agar. The number of colony forming units (CFUs) was recorded after 5 days of incubation at 30°C (CFU method). The experiment was performed twice. Results were expressed as mean percent culturability.

To assess the ability of 8D-HYPOx cells to grow in liquid medium, we determined the probability for a cell to grow by seeding, for each condition, 576 wells (six 96-well plates) with 100 yeasts/well and counting the number of positive wells at the end of the incubation time, taking into account the Poisson’s law (plate method). Plates were incubated 5 days at 30°C in an Infors HT^®^ multitron pro at 150 rpm. Two independent experiments were performed for each condition. The probability (p) of a cell to grow was calculated using the following formula, where X represents the number of positive wells, Y the number of wells seeded.

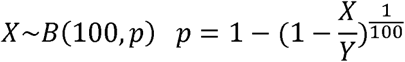

### Phagocytosis assay

Interaction with macrophages was performed as described previously [92] using the J774 cell line, the monoclonal anti-capsular polysaccharide antibody E1 as an opsonin [93], calcofluor-stained yeasts and a yeasts: macrophages ratio of 5:1. Phagocytosis was assessed by flow cytometry using the Guava easyCyte 12HT Benchtop Flow Cytometer^®^ (Merck).

### Phenotypic microarrays

Using phenotypic microarray technology (Biolog^®^, Hayward,CA), we analyzed the metabolic capacities of yeasts incubated VBNC/8D-NORMOx (both with biological duplicates), using as controls either STAT or dead cells (8 days in hypoxia in a YPD pH = 5) (only 3 conditions, i.e. VBNC, 8D-NORMOx and control could be tested in each series). The Biolog^®^ apparatus allows quantitative measurement of the metabolism over time based on the physiological state of the yeast cells independently of cell division and based on mitochondrial activity (proton and electron generation) [94]. Reduction of tetrazolium dye that absorbed charge produced by cell respiration results in a purple color change that is recorded over time (Omnilog unit, equivalent of the Optical Density measured at 590nm). The optical density measured in the well provide information on the intensity of the corresponding metabolic activity. The effect of carbon, nitrogen, phosphorus, and sulfur sources on the physiological state of the cell and cell respiration was examined in response to various osmotic conditions and to different pHs (plates PM1, PM2A, PM3B, PM4A, PM6, PM7 and PM8). A tetrazolium dye reacting to the redox status of the well turns purple when respiration takes place which has been shown to be a good proxy for the ability of strains to grow on a substrate [95]. Phenotype microarray was done according to instructions from the manufacturer (Biolog, Hayward, CA) with the following exceptions. Briefly, ≈7×10^8^ cells were washed 3 times in PBS and then resuspend in 27 mL of PBS. For each condition, the analysis was performed based on the results obtained after 6 days of incubation. Overall, 761 compounds were tested and 240 compounds that were positive in the dead cells condition were excluded from the analysis. The heatmap and clustering of the 521 remaining conditions were performed using R studio v1.1.456 with ggplot package using the OD values measured by the Biolog^®^ normalized between 0 and 1 using the following formulae *zi*=(*xi*−min(*x*)) / (max(*x*)−min(*x*)). For the endpoint comparison, only positive wells in each condition were compared (393 in STAT, 279 in VBNC duplicate 1, 200 in VBNC duplicate 2, 454 in 8D-NORMOx duplicate 1 and 380 in 8D-NORMOx duplicate 2).

### Mitochondrial nucleic acid quantification

Cells suspensions corresponding to STAT or VBNC (1 mL each in biological triplicates) were centrifuged at 4 000rpm for 5 min at 4°C before a flash freeze step in −80°C ethanol solution and storage at −80°C before extraction. The RNA extraction protocol was adapted from [96]. Briefly, samples were lyophilized during 18h. RNAs were extracted in 3 successive steps: i) extraction performed in Trizol reagent (Invitrogen TriReagent (Sigma-Aldrich, St-Louis MO 63103 USA)), ii) “separation” by adding chloroform (Invitrogen) iii) overnight isopropanol precipitation at −20°C.

The DNA protocol extraction was a manual method based on 3 steps. Briefly, a first stage consisted in lysing the cell pellets by beat beating with MagNA Lyser Instrument (Roche Diagnostics; 2 rounds at 7 000 r/min for 45 s each) in a solution of 2% Triton X-100 (Sigma) and 1% of SDS (Biosolve BV Chimie, 5555 CE Valkenswaard, The Netherlands), NaCl 100mM; Tris 10mM pH 8.0; EDTA 1mM. Then, an extraction step in phenol/chloroform solution (Phenol:Chloroform:isoamyl alcohol pH 8.0, 1 mM EDA (Sigma-Aldrich, St-Louis MO 63103 USA) and a precipitation step in absolute ethanol and 5M of ammonium acetate solution were done. RNAs and DNAs were quantified using the Nanodrop Spectophotometer (ThermoFisher, Scientific, Inc.). All extracts were stored at −80°C before analysis.

### qPCR and reverse transcriptase qPCR

Briefly, mtLSU primers were designed with Primers. 3 vs 4.1.0, while all other specific primers were designed to be compatible with the Human Universal Probe Library set (90 probes, octamer, Roche Diagnostics) (Supplementary Table 9) [23]. All reverse transcriptase qPCR assays for a given target were performed simultaneously for all samples using the LightCycler Multiplex RNA Virus Master Kit (Roche diagnostics). Each sample was normalized with the geometric mean of the quantification cycle (Cq) of the corresponding GAPDH and ACT housekeeping genes expression [97] and fold changes calculated according to Pfaffl after determining the efficiency of each PCR assay [98].

### Secretome and proteome analysis

Three biological replicates were performed kinetically upon incubation D4, D6 and D8 in hypoxia and normoxia starting from step3 (Supplementary Figure 1) corresponding to stationary phase (STAT). The suspensions were centrifuged at 4000rpm and the supernatants (secretome) filtrated through a 0.22 µM membrane (Millex GV, Merck Millipore, Burlington, USA). The corresponding cell pellets (proteome) were extracted in parallel. For each biological replicate, three technical replicates were pooled to increase the amount of recovered proteins for the supernatants.

All samples (supernatant or pellet) were lyophilized overnight after flashed freezing in liquid nitrogen. We then adapted a protocol established to purify a sample rich in carbohydrates [99]. The lyophilized samples were resuspended in 0.5 mL of sterile milliQ water and the following 3 steps-protocol was done: i) the samples were precipitated at 4°C by addition of 1 mL of 10% trichloroacetic acid (TCA) in acetone (kept at-20°C) followed by incubation for 5 m. After centrifugation at 20000g for 10 minutes at −20°C, the pellet was resuspended in 100% acetone with 10mM dithiothreitol (DTT), washed twice in this solution, and air dried; ii) the pellet was then solubilized in sodium dodecyl sulfate (SDS) extraction buffer containing 1% SDS, 0.15 M Tris HCl pH=8.8, 0.1M DTT, 1 mM ethylene diamine tetra acetic acid (EDTA), 2 mM phenylmethylsulfonyl fluoride (PMSF), and incubated for 1 h at room temperature and vortexed every 20 m. A 15000g centrifugation for 10 min at room temperature allowed recovery of the supernatant; iii) Phenol was added at equal volume to the supernatant and the tube was vortexed for 3 m. After centrifugation at 20000 g for 5 min at room temperature, the denser phenolic phase was recovered in the lower part. An equal volume of washing solution (Tris HCl pH 8.0, 10 mM, 1 mM EDTA, 0.7 M sucrose) was added and the tube vortexed for 3 m. The phenolic phase was then recovered on the upper part and transferred to a new tube. Then, 0.1M ammonium acetate in methanol (−20°C) was added and the mixture stored at −20 ° C for 30 m. The precipitate was pelleted by 20000g centrifugation for 10 min at −20°C, and washed first with 0.1M ammonium acetate in methanol and then with 100% acetone. After a last centrifugation step (20000 g for 10 min at −20°C), the pellet was air-dried and resuspended in 350 μl of denaturing buffer (8M urea, 100 mM TrisHCl pH 8.0). Two μL of each sample were then used for dosage with the Biorad DC^®^ dosing kit according to the manufacturer’s recommendations. The remaining solution was stored at −80°C until analysis.

### High pressure liquid chromatography coupled to tandem mass spectrometry (nano HPLC MS/MS)

In solution protein digestion. Samples were re-suspended in 350 µL 8 M urea/ 100 mM Tris HCl pH 8.5 after TCA/acetone precipitation and phenol extraction. Briefly, samples were reduced with 5 mM tris(2-carboxyethyl)phosphine (TCEP) for 30 min at room temperature and alkylated with 10 mM iodoacetamide for 30 min at room temperature in the dark. Then, proteins were firstly digested for 5 h at 37°C with 250 ng rLys-C Mass Spec Grade (Promega, Madison, USA). Samples were then diluted 4-fold with 100 mM Tris HCl pH 8.5 to reach a concentration of 2M urea and then re-incubated overnight at 37°C with 1 µg Sequencing Grade Modified Trypsin (Promega, Madison, WI, USA). A second incubation with the same amount of trypsin (5 h at 37°C) was performed to ensure a complete digestion. Digestion was stopped by adding formic acid to 5 % final concentration and peptides were desalted and concentrated on Sep-Pak C_18_ SPE cartridge (Waters, Milford, MA, USA) according to manufacturer’s instructions.

Mass spectrometry data analysis. Tryptic peptides were analyzed on a Q Exactive Plus instrument (Thermo Fisher Scientific, Bremen) coupled with an EASY nLC 1000 chromatography system (Thermo Fisher Scientific). Sample was loaded on an in-house packed 30 cm nano-HPLC column (75 μm inner diameter) with C18 resin (1.9 μm particles, 100 Å pore size, Reprosil-Pur Basic C18-HD resin, Dr. Maisch GmbH, Ammerbuch-Entringen, Germany) and equilibrated in 98 % solvent A (H2O, 0.1 % FA) and 2 % solvent B (ACN, 0.1 % FA). Peptides were first eluted using a 2 to 18 % gradient of solvent B during 112 m, then a 18 to 30 % gradient of solvent B during 35 m, a 30 to 45 % gradient of solvent B during 15 min and finally a 45 to 60 % gradient of solvent B during 5 min all at 250 nL.m^-1^ flow rate. The instrument method for the Q Exactive Plus was set up in DDA mode (Data Dependent Acquisition). After a survey scan in the Orbitrap (resolution 70 000), the 10 most intense precursor ions were selected for HCD fragmentation with a normalized collision energy set up to 27. Charge state screening was enabled, and precursors with unknown charge state or a charge state of 1 and >7 were excluded. Dynamic exclusion was enabled for 45 s.

Data processing and analysis. All data were searched using Andromeda [100] with MaxQuant software [101,102] version 1.5.3.8 against the H99 strain proteins database (6980 entries) and the H99 strain mitochondria proteins database (13 entries) concatenated with usual known mass spectrometry contaminants and reversed sequences of all entries. Andromeda searches were performed choosing trypsin as specific enzyme with a maximum number of two missed cleavages. Possible modifications included carbamidomethylation (Cys, fixed), oxidation (Met, variable) and Nter acetylation (variable). The mass tolerance in MS was set to 20 ppm for the first search then 6 ppm for the main search and 10 ppm for the MS/MS. Maximum peptide charge was set to seven and five amino acids were required as minimum peptide length. The “match between runs” feature was applied for all samples with a maximal retention time window of 2 m. One unique peptide to the protein group was required for the protein identification. A false discovery rate (FDR) cutoff of 1 % was applied at the peptide and protein levels.

MaxLFQ, Maxquant’s label-free quantification (LFQ) algorithm was used to calculate protein intensity profiles across samples [103]. An LFQ value was assigned when the protein had at least 2 peptides, of which 1 was unique. The median reproducibility by samples for secretomic was 87.1% [82.5-90.6] and 86.5% [84.1-88.9] for cellular proteins.

Statistical analysis of differentially expressed proteins. Proteins for which we had no value for a given condition and time were removed. Remaining missing values were imputed using the structured least Squares algorithm (package imp4p) [104].

To get a global overview of the data, a principal coordinate analysis (PCoA) or a principal component analysis (PCA) was used [105]. The differential analysis was done with the linear model approach proposed by the package R limma (v3.30.13) [106]. We defined a model that included condition (hypoxia or normoxia), time (D0=STAT, D4, D6, and D8) and replicate (1 to 3) variables as main effects and interactions between conditions and times as well as conditions and replicates. The latter interaction was useful to model the pairing between measurements corresponding to the same condition. The proteins differentially expressed over time are selected according to the following criteria: change with respect to the initial value of log2 and a p <0.05. The p-values obtained were adjusted by the method of Benjamini-Hochberg [107].

### Analysis of enriched processes for secreted proteins “Gene Ontology (GO) enrichment analysis”

The fungifun2 v2.2.8 BETA [108] site (https://elbe.hki-jena.de/fungifun/fungifun.php) was used by selecting for each sample of secreted proteins the corresponding cellular protein background and to study the cellular proteins the background was the whole database. For this analysis we selected only the proteins identified in the 3 replicates per condition. For each analysis, the following options were selected: p value <0.01, hypergeometric distribution test, over-representation, p-read corrected by Benjamini-Hochberg, only directly annotated association. Hypothetical proteins, which constitute 20-30% of the annotated proteins, were not represented.

### Venn Diagram

Venn diagram was generated using Jvenn [109] (http://jvenn.toulouse.inra.fr/app/index.html) with proteins only present in the 3 replicates for each condition (secretome and proteome).

### Protein/protein interactions analysis

Analysis of protein-protein interactions of intracellular proteins regulated during hypoxia. STRING v10.5 software (https://string-db.org) was used [110]. As this software used the strain JEC21 annotation, the list of orthologous proteins was transferred from H99 to JEC21 with a 62/63 protein match. The nodes directly related to the input were indicated with colors while the nodes of a higher iteration/depth were indicated in white.

### KEGG pathway mapping and fatty acid degradation cycle reconstruction

KEGG pathway database (https://www.genome.jp/kegg-bin) was used to map the pathways. It was rendered with the intracellular proteins significantly and differently produced in hypoxia: 63 proteins were transferred from H99 to JEC21. Nineteen proteins belonged to general metabolic pathway and more precisely 8 to fatty acid degradation (“map” pathways were not colored, *Cryptococcus*-specific pathways were colored green, entries were colored red). Another fatty acid degradation cycle was reconstructed from yeast genome database cycle (https://www.yeastgenome.org/) and the corresponding proteins were found in *Cryptococcus neoformans* H99 genome using BLAST (https://blast.ncbi.nlm.nih.gov/Blast.cgi).

### RNAseq analysis

For transcriptome analysis, yeasts incubated in 8D-HYPOx/8D-NORMOx, as well as STAT and logarithmic (LOG) phase yeasts were studied in triplicates for two different sets, one containing spiked *Saccharomyces cerevisiae* organisms (w_spike) and one with *C. neoformans* cells only (w/o_spike). The former strategy was used to obtain an internal standard of the level of expression of the whole transcripts and adjust the level of expression of each gene to that of the constant expression of *S. cerevisiae*. In each spiked sample, 15% of LOG phase *S. cerevisiae* cells were added to *C. neoformans* cells. Triplicates of yeasts suspensions in the different conditions (total of 2.55×10^8^ *C. neoformans* cells) were spiked (w_spike) or not (w/o_spike) with 4.5×10^7^ cells (representing then 15% of the cells) of *S. cerevisiae* cells (culture in exponential phase). All samples were then processed as described above in the Mitochondrial nucleic acid quantification section.

We used 1 μg of total RNA to purify polyadenylated mRNAs and to build an RNA library, using TruSeq Stranded mRNA Sample Prep Kit (Illumina, #RS-122-9004DOC) as recommended by the manufacturer. Directional library was checked for concentration and quality on RNA chips with the Bioanalyzer Agilent. More precise and accurate quantification was performed with sensitive fluorescent-based quantitation assays (“Quant-It” assays kit and QuBit fluorometer, Invitrogen). Samples were normalized at 2 nM concentrations, multiplexed 4 samples per lane and then denatured at a concentration of 1 nM using 0.1 M NaOH at room temperature. Each sample was finally loaded on the flowcell at 9 pM. Sequencing of the 24 samples was performed on the HiSeq 2500 sequencer (Illumina) in 65 bases single-end mode.

### QC, mapping, counting, and global transcriptome level analysis

Reads were cleaned of adapter sequences and low-quality sequences using cutadapt version 1.11 (Marcel Martin. Cutadapt removes adapter sequences from high-throughput sequencing reads [111]). Only sequences at least 25 nucleotides in length were considered for further analysis. STAR version 2.5.0a [112], with default parameters, was used for alignment on the reference genome (PRJNA411 from NCBI Bioproject). Read counts per gene were measured using featureCounts version 1.4.6-p3 [113] from Subreads package (parameters: -t gene -s 0). Results are summarized using multiqc version 0.7 [114]. Count data were analyzed using R version 3.3.1 and the Bioconductor package DESeq2 version 1.12.3 [115]. Normalization and dispersion estimations were performed with DESeq2 using the default parameters but statistical tests for differential expression were performed without applying the independent filtering algorithm.

For w/o_spike samples, a generalized linear model was set in order to test for the differential expression between STAT, 8D-HYPOx/8D-NORMOx and LOG conditions. For each pairwise comparison, raw p-values were adjusted for multiple testing according to the Benjamini and Hochberg (BH) procedure and genes with an adjusted p-value lower than 0.01 were considered differentially expressed.

For w_spike samples, mean normalized counts were calculated and the rank of each gene determined for each gene in each condition. High and low rank values corresponded to genes with the highest and lowest expression relative to spiked RNA, respectively.

### Differential expression analysis

To identify the expression differences across conditions, the read counts were filtered and normalized using the R package “DESeq2”. Genes with read counts below a threshold of 10 in all samples were filtered out. After normalization of read counts, differentially expressed genes for each condition were identified using the generalized linear model, which perform pairwise comparisons among each of the conditions (STAT, 8D-HYPOx, 8D-NORMOx and LOG). Genes passing the threshold, (FDR <5%), were considered significantly differentially expressed and were then considered to be up or down-regulated according to the direction of the fold change. The k-means clustering analyses were performed using R tools, “kmeans” and “heatmap.2”. To discover the underlying pathways of these differentially expressed genes, we also carried out functional annotation analysis by summarizing the data, including known H99 gene annotation, *S. cerevisiae* gene ortholog, Gene Ontology (GO), KEGG pathway, and pfam domain, from FungiDB database [116]. GO enrichment and KEGG pathway enrichment for these genes were then assessed by using Fisher’s Exact Test with multiple testing correction of FDR threshold at 5%. Furthermore, conserved domains of each *C. neoformans* gene were assessed by using the amino acid sequence to search the NCBI Conserved Domain Database (CDD) [117]. The presence of signal peptide sites was predicted by SignalP 4.1 [118].

### Statistical analysis

The statistical tests used for the comparisons between the different conditions were t-test, Mann Whitney, Wilcoxon tests according to the distribution of the data.

### Data access

All RNA sequence data from this study have been submitted to NCBI (https://www.ncbi.nlm.nih.gov/geo) under accession number GSE118549.

The mass spectrometry proteomics data have been deposited to the ProteomeXchange Consortium via the PRIDE partner repository with the dataset identifier PXD012570 (https://cran.r-project.org/).

## Supplementary figures legends

**Supplementary figure 1: *in vitro* protocol for dormant cell generation.** The protocol followed several steps: (1) *Cryptocococcus neoformans* cells of a stock culture frozen at −80°C were grown on Sabouraud agar plate for 2 to 5 days at room temperature; (2) 10^7^ cells were then suspended in 10 mL yeast extract peptone dextrose 2% (YPD) and incubated with lateral shaking (150 rpm) at 30°C for 22 hours (stationary phase-STAT); (3) 100 μl, 2×10^7^ cells were incubated again until the STAT and then placed in hypoxia or normoxia in the dark at 30°C (4) for up to 8 days when the resultant cells in hypoxia and normoxia were analyzed.

**Supplementary figure 2: Cell body size, capsule and cell wall thickness. A.** Median cell sizes were similar in 8D-HYPOx, 8D-NORMOx and STAT. **B.** Median capsule size was lower in 8D-HYPOx, 8D-NORMOx compared to STAT (*p<0.01, 100 cells measured). **C.** Cell wall was thicker in 8D-HYPOx compared to STAT (*p=0.0152, 50 cells measured).

**Supplementary figure 3: Flow cytometry diagrams assessing capsular, vacuolar structure and viability in STAT, 8D-NORMOx and 8D-HYPOx. A.** Histograms of fluorescence intensity showing no difference in binding pattern after staining with the E1 anti-glucuronoxylomannan monoclonal antibody **B.** More intense vacuolar staining with MDY-64 in 8D-HYPOx, 8D-NORMOx conditions (one representative experiment out of the 3 independent experiments performed is shown) **C.** Viability was assessed by membrane permeability staining (LIVE/DEAD, LVD) showing almost 99% of live cells in STAT cells and 100% of dead cells in heat-killed cells (Left panel). Plasma membrane was intact for more (87.2% [83.2-88.6]) cells in 8D-HYPOx, than in 8D-NORMOx (53.8% [50.9-60.5]). Experiments were done in triplicate and a representative diagram is shown.

**Supplementary figure 4: Latency was influenced by medium and cell concentration**

Growth of STAT and 8D-HYPOx cells was assessed using the BioScreen^®^ apparatus. Serial dilutions of 8D-HYPOx and STAT cells increased the latency in YPD (**A**) and in MM (**B**). Latency curves extrapolations showed for both STAT and 8D-HYPOx cells, global latency was decreased in YPD compared to MM. Each point represents the median ± IQR of the latency of 3 independent experiments. **C.** Experimental set up used for the determination of the probability of growth per cell (culturability). Hundred yeasts cells per well were plated in 96-well plates in each condition (MM at 10 or 100% ± pantothenic acid (PA) at 125µM). The number of positive wells per plate were determined and the probability of growth per plate was calculated (see M&M section) D. 8D-HYPOx were exposed to macrophages during two hours in the presence of opsonin. Culturability was similar in phagocytosed 8D-HYPOx cells compared to controls. Each dot represents the calculated culturability. Two independents experiments are pooled.

**Supplementary figure 5: ROS and RNS production were decreased in VBNC**

Testing the ROS and RNS in VBNC vs STAT cells found a slight decreased in VBNC. ROS (left panel) and RNS (right panel) productions were measured using fluorescence probes and were significantly lower in VBNC compared to STAT cells respectively (*p<0.01). As expected, the addition of H_2_0_2_ as a positive control increased ROS and RNS levels.

**Supplementary figure 6: The number of identified proteins evolved differently between secreted and cellular proteins in VBNC and 8D-NORMOx**. The number of secreted protein number significantly decreased over time in VBNC (* p<0.01) (**A**) while the protein concentration tended to increase in 8D-NORMOx (**B**). The number of cellular proteins remained stable in both conditions (**C**) while the protein concentration tended to increase (**D**). The experiments were done in triplicates (median±IQR].

**Supplementary figure 7: Venn diagrams of STAT, VBNC and 8D-NORMOx for secreted and cellular proteins. A.** A hundred and seven cellular proteins were only present in hypoxia and 2408 were common to the 3 experimental conditions. B. Seventeen secreted proteins were only present in hypoxia and 252 were common to the 3 experimental conditions.

**Supplementary figure 8: GO enrichment analysis for molecular function (A,B) and cellular component (C,D) showed a particular pattern in hypoxia for secreted proteins (A,C) but not for cellular (B,D) proteins. A.** The major enriched molecular function process for secreted proteins in hypoxia was structural constituents of ribosome and in normoxia catalytic and oxidoreductase activity. **B.** For cellular component, the major enriched cellular components were the ribosome, ribonucleoprotein complex and intracellular in hypoxia and mainly ribosome in normoxia. **C.** The major enriched molecular function process for cellular proteins in hypoxia were the same as normoxia: nucleotide binding, transferase activity and catalytic activity. **D.** For cellular component, the major enriched cellular component for both hypoxia and normoxia were cytoplasm, ribosome, ribonucleoprotein complex and intracellular.

## Acknowledgements

We would like to acknowledge Christina A. Cuomo, Laurent Châtre, Guilhem Janbon, Jean-Yves Coppée, Pierre Rocheteau, Pierre Henri Commere, Christine Schmitt, Olivier Gorgette, Jacomine Krijnse-Locker, Caroline Proux for their comments on this work and help at different steps of the study.

---

**Supporting Table 1:** Optical density values for the 10 Biolog^®^ plates.

**Supporting Table 2:** Number and concentration of secreted or cellular proteins

**Supporting Table 3:** List of proteins used to build the Venn diagram

**Supporting Table 4:** List of secreted and cellular protein found in each and combined conditions

**Supporting Table 5:** Proportion of biological processes, Molecular functions and cellar components enriched in secreted and cellular proteins for each condition

**Supporting Table 6:** Regulated secreted and cellular proteins in normoxia and hypoxia compared to STAT

**Supporting Table 7:** List of GO and CNAG transcripts enriched in each k-mean cluster obtained from the transcriptome analysis

**Supporting Table 8:** Strains used in this study

**Supporting Table 9:** Primers used in this study

