## Supplementary figures and images for "*Cryptococcus neoformans* resists to drastic conditions by switching to viable but non-culturable cell phenotype" (from another version of this preprint: version 1, 2019-02-18)

### Supplemental Figure 1

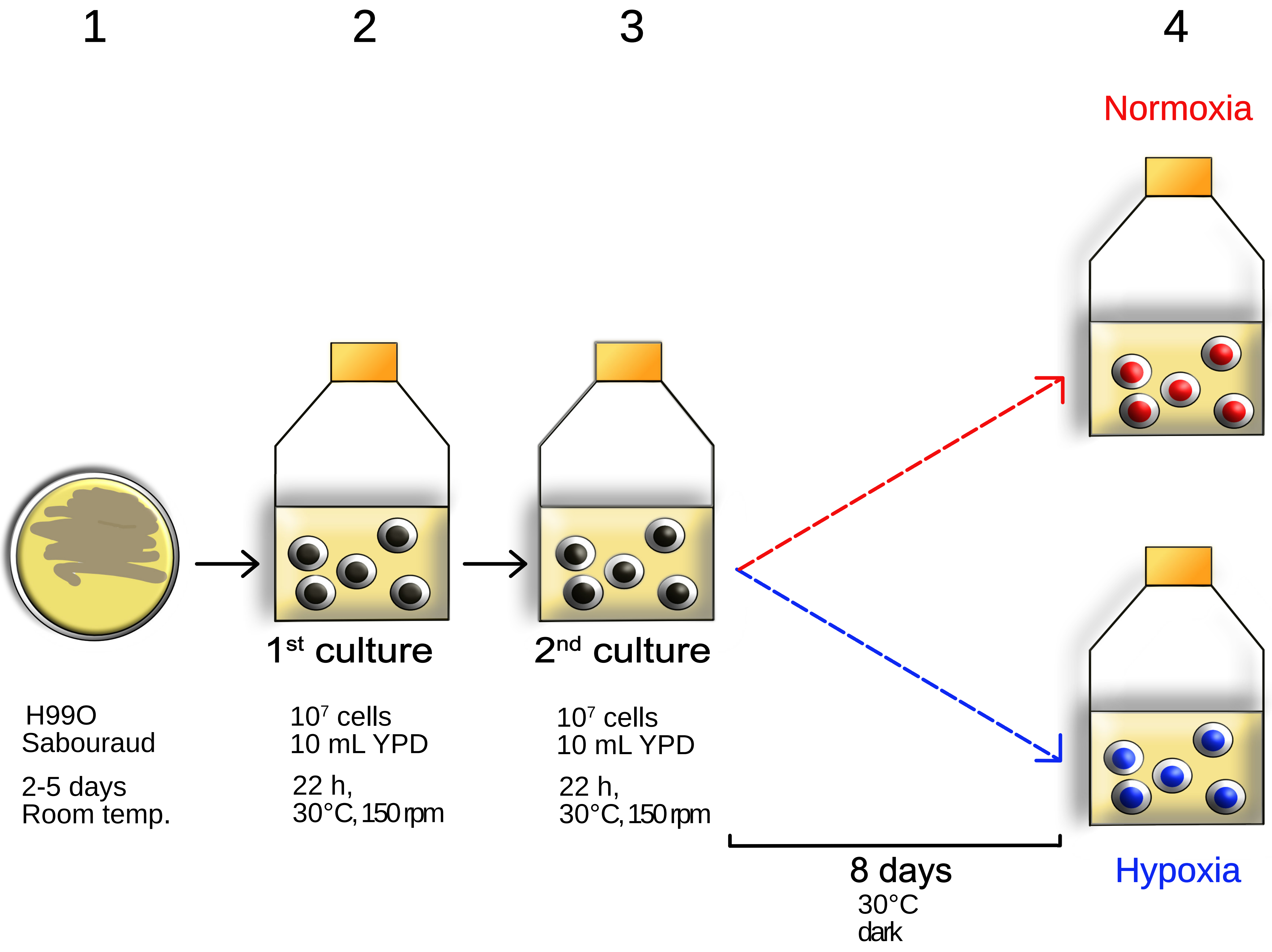

### Supplemental Figure 2

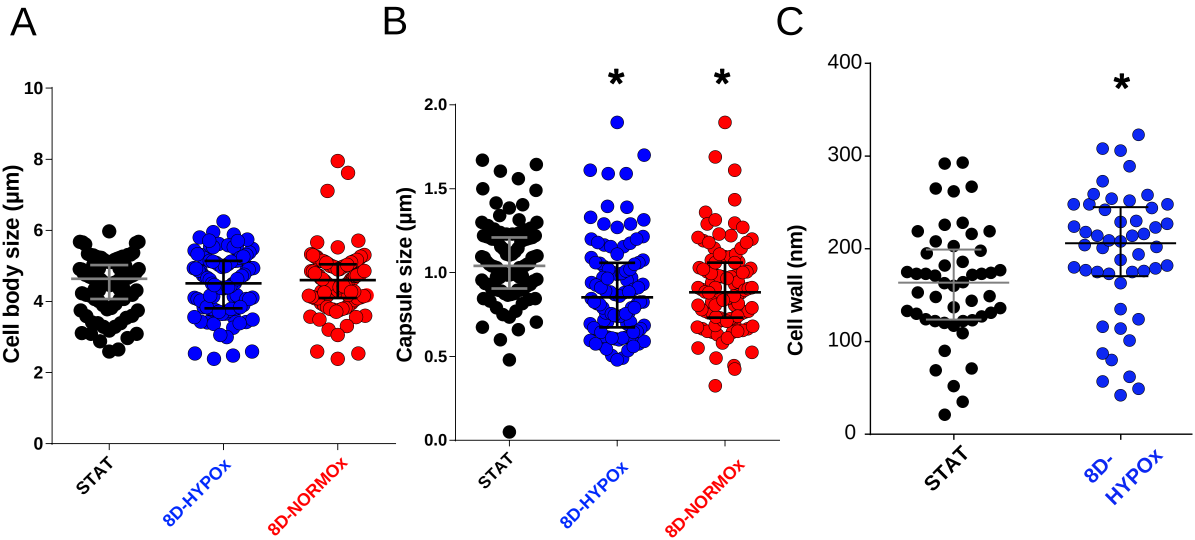

### Supplemental Figure 3

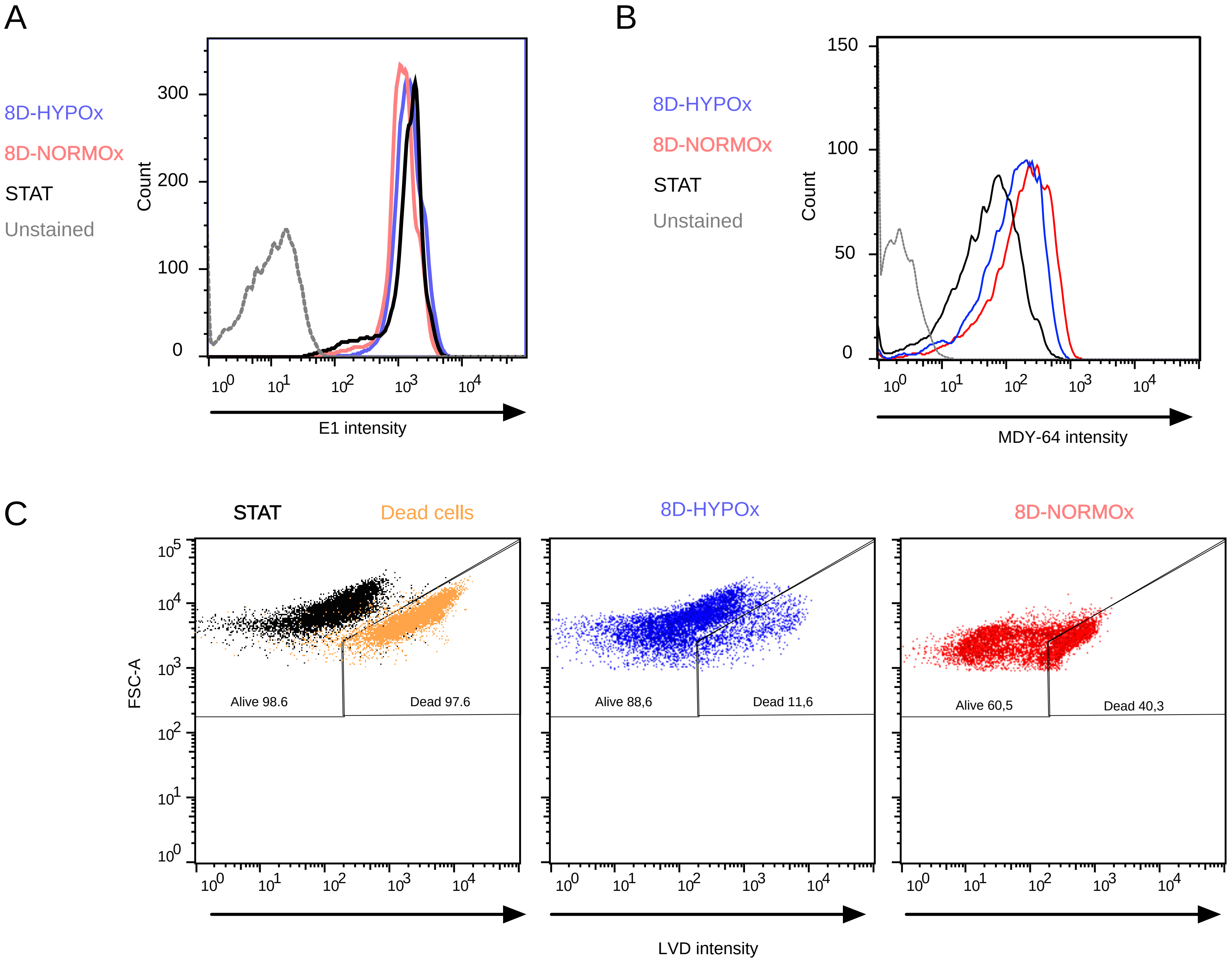

### Supplemental Figure 4

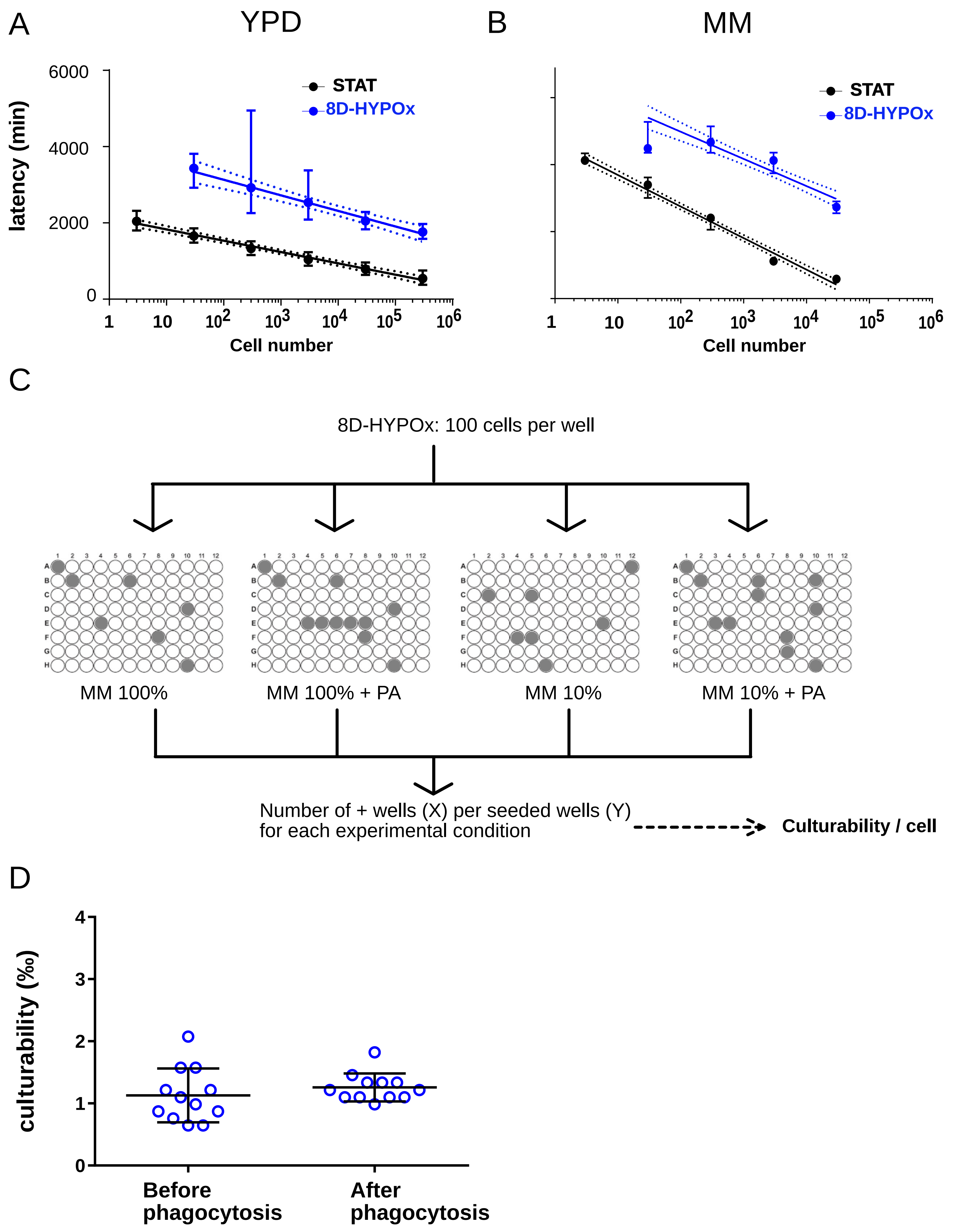

### Supplemental Figure 5

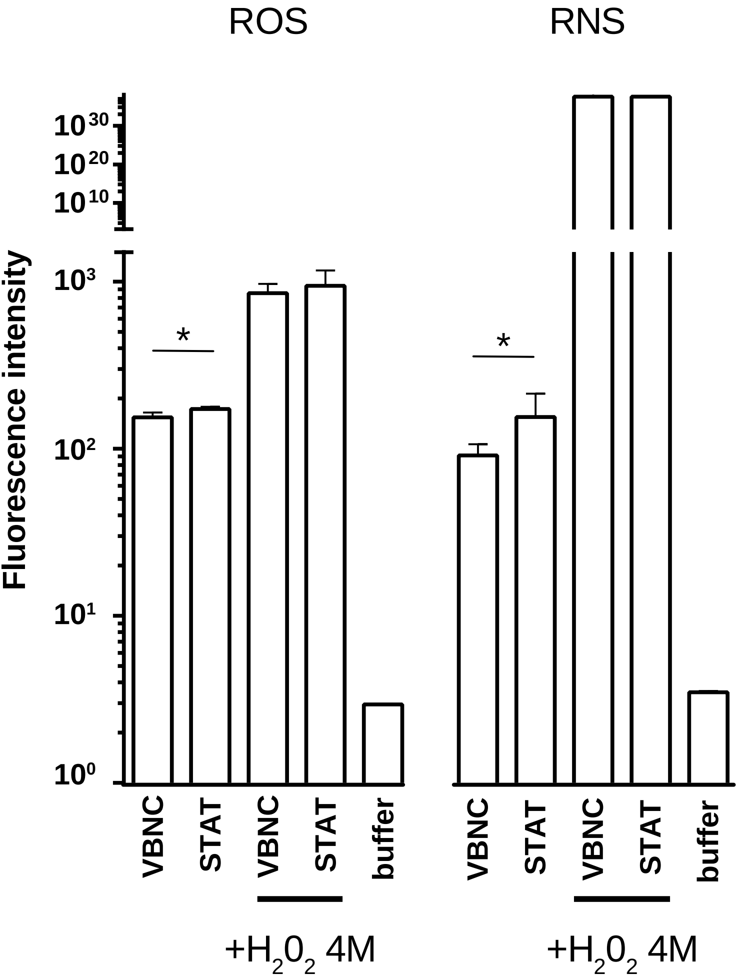

### Supplemental Figure 6

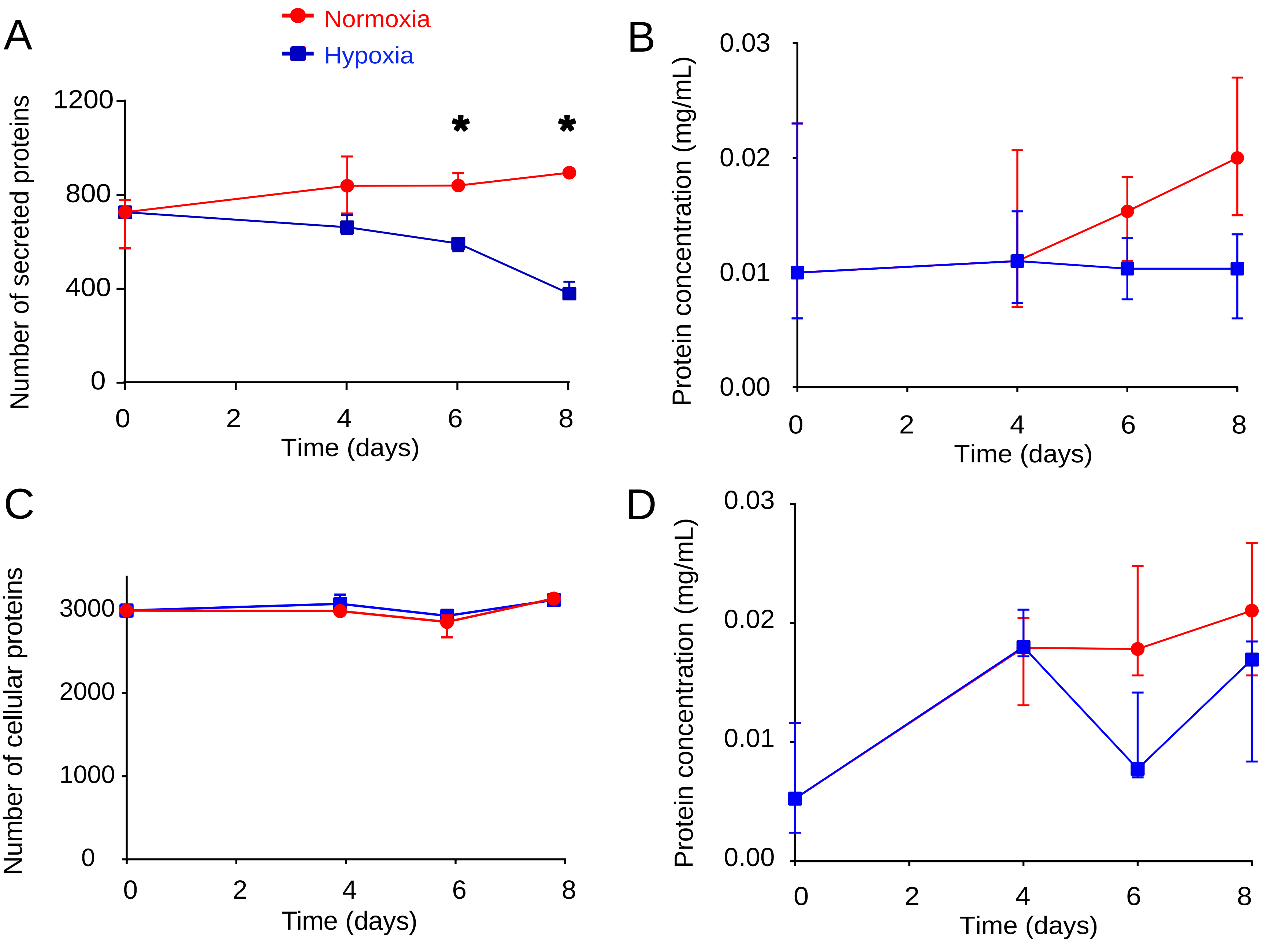

### Supplemental Figure 7

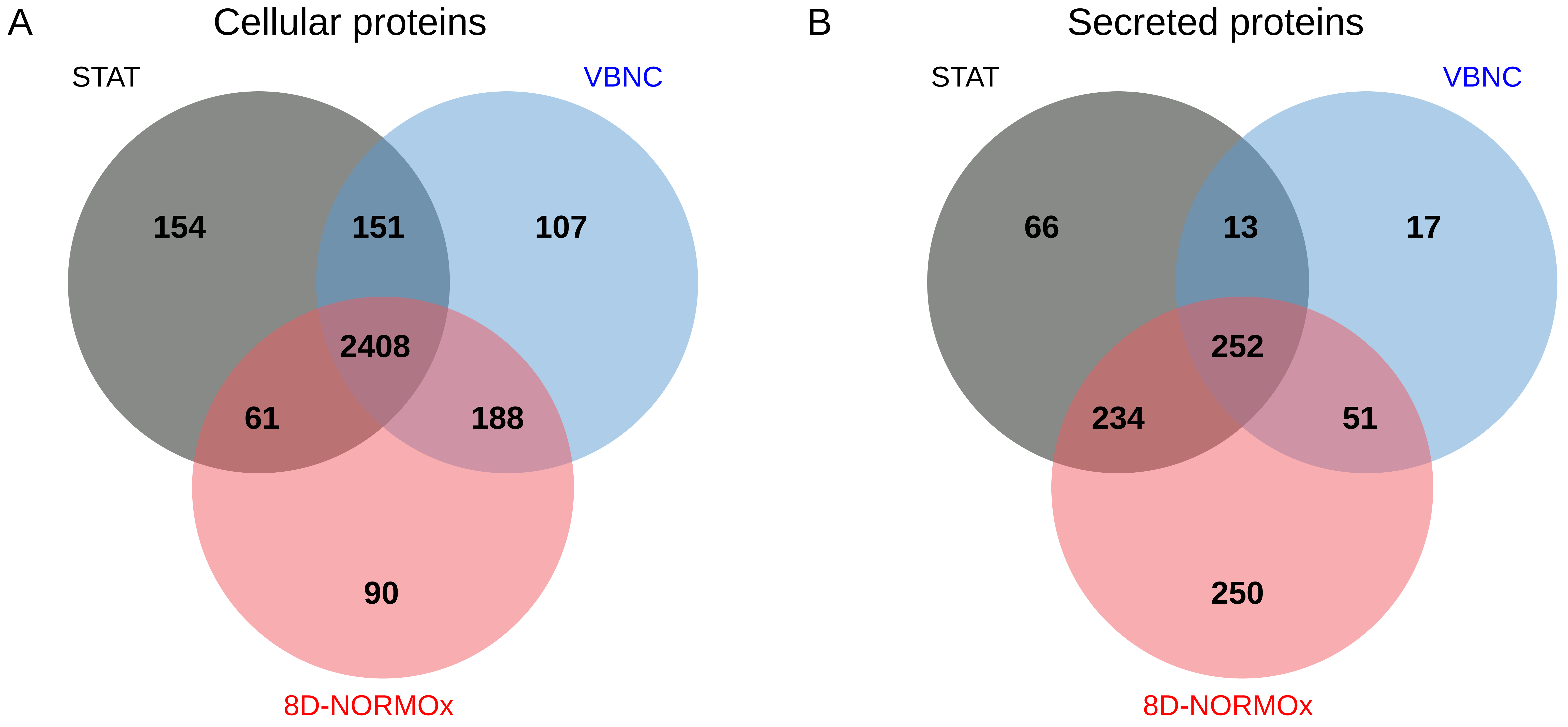

### Supplemental Figure 8

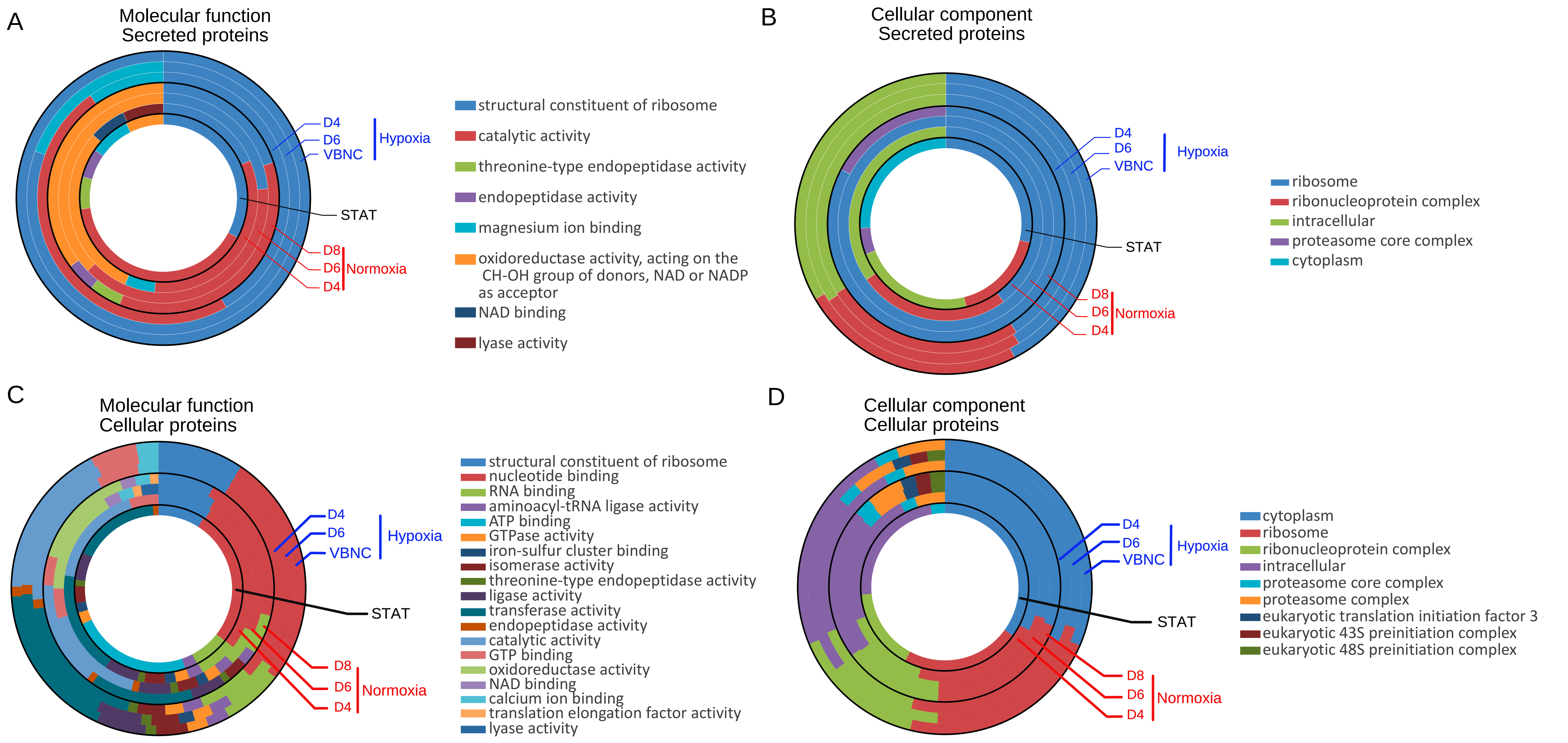
