## Supplemental Table 8 for "*Cryptococcus neoformans* resists to drastic conditions by switching to viable but non-culturable cell phenotype" (from another version of this preprint: version 1, 2019-02-18)

**Table S8: Strains used in this study**

| Strain denomination | Reference or Source |
| --- | --- |
| H99O | J. Heitman (Duke University, Durham, NC, USA |
| KN99α | Madhani collection |
| *CNAG_02489Δ* KN99α | Madhani collection |
| *CNAG_04688Δ* KN99α | Madhani collection |
| *CNAG_06628Δ* KN99α | Madhani collection |
| *CNAG_03019Δ* KN99α | Madhani collection |
| *CNAG_00490Δ* KN99α | Madhani collection |
| *CNAG_07747Δ* KN99α | Madhani collection |
| *CNAG_04531Δ* KN99α | Madhani collection |
| *CNAG_00524Δ* KN99α | Madhani collection |
| *CNAG_05721Δ* KN99α | Madhani collection |
